# Museum collections and machine learning guide discovery of novel coronaviruses and paramyxoviruses

**DOI:** 10.1101/2025.09.11.675601

**Authors:** Maya M. Juman, Molly M. McDonough, Adam W. Ferguson, Barbara A. Han, Frank Bapeamoni Andemwana, Bertin Murhabale Cisirika, Charles Kahindo, Luis M.P. Ceríaco, Steven M. Goodman, Bruce D. Patterson, Greg F. Albery, Colin J. Carlson, Daniel J. Becker

**Affiliations:** Department of Veterinary Medicine, University of Cambridge, Cambridge, UK; Department of Biological Sciences, Chicago State University, Chicago, IL, USA; Negaunee Integrative Research Center, Field Museum of Natural History, Chicago, IL, USA; Gantz Family Collections Center, Field Museum of Natural History, Chicago, IL, USA; Cary Institute of Ecosystem Studies, Millbrook, NY, USA; Faculté des Sciences, Université de Kisangani, Kisangani, DRC; Département de Biologie, Faculté des Sciences, Université Officielle de Bukavu, Bukavu, DRC; Centro de Investigação em Biodiversidade e Recursos Genéticos, Universidade do Porto, Portugal; BIOPOLIS Program in Genomics, Biodiversity and Land Planning, Portugal; School of Natural Sciences, Trinity College Dublin, Dublin, Ireland; Department of Epidemiology of Microbial Diseases, Yale University, New Haven, CT, USA; School of Biological Sciences, University of Oklahoma, Norman, OK, USA

## Abstract

Natural history museum collections are valuable but underutilized resources for viral discovery, offering opportunities to test hypotheses about viral occurrence across space, time, and taxonomic groups. We developed machine learning models of bat host suitability to guide coronavirus and paramyxovirus screening of 1330 and 491 tissues, respectively, in a museum collection. For the first time, we recovered coronavirus (*n* = 16) and paramyxovirus (*n* = 3) sequences from archived museum tissues, confirming three novel coronavirus host species and three novel paramyxovirus host species (3% and 33% prediction success rate, respectively). These sequences included a SARS-like coronavirus and an orthoparamyxovirus from Angolan *Rhinolophus fumigatus* specimens collected in June 2019, suggesting that viruses with epidemic potential may be more widespread in sub-Saharan Africa than previously believed. Our study demonstrates the value of combining predictive modeling and collections-based viral discovery, particularly for filling outstanding sampling gaps and investigating changes in host–virus associations over time.

## Introduction

Most emerging human infectious diseases are zoonotic, and recent anthropogenic changes are driving increased spillover from and between wildlife reservoir hosts (Eby et al. 2023; Gibb et al. 2024; Carlson et al. 2025). Predicting and mitigating spillover relies on pathogen discovery, host identification, and long-term study of transmission dynamics; however, while crucial, these studies are costly and logistically difficult to conduct in the field, and they often rely on retrospective wildlife sampling during and after outbreaks (Kuiken et al. 2005; Plowright, Becker, McCallum, et al. 2019). A promising—but underutilised—complementary approach involves the screening of archived samples, particularly natural history museum specimens (Harmon et al. 2019; Thompson et al. 2021).

The use of existing natural history collections for retroactive pathogen surveillance dates back to at least the 1990s, when decades-old specimens were used to identify the reservoir host of a novel hantavirus (Yates et al. 2002). Recent sampling efforts have uncovered DNA viruses and tick-borne RNA viruses in mammal specimens (Lund et al. 2023; Hämmerle et al. 2024; Gieraths et al. 2025). Other pathogens can also be identified in museum collections, including bacteria (Leber et al. 2022), fungi (Carvalho et al. 2017), protozoa (Galen et al. 2019), and helminths (Preisser et al. 2022). A shift toward the “extended specimen” model of collections has encouraged the extraction and digitization of all associated biological material and metadata. Thus, specimens serve as vouchers of host– pathogen interactions that can illuminate and even expand the spatial, temporal, and taxonomic ranges of currently known associations. These data can also facilitate investigations of shifts in pathogen abundance, diversity, and distribution in response to anthropogenic change (Wood et al. 2023). Screening specimens is cost-effective, efficient, and proactive relative to traditional, responsive field-based sampling (Colella et al. 2021). Still, despite their numerous benefits, museum collections remain largely untapped resources for pathogen discovery and putative reservoir host identification.

Globally, natural history museums house over one billion specimens, which in theory could enable myriad investigations of pathogens across space and time (Johnson et al. 2023). However, the vastness of these collections also poses challenges, particularly for exploratory studies of emerging pathogens where host species are largely or entirely unknown. Predictive modeling has proven effective in narrowing down the list of putative hosts by identifying unsampled species or clades that are likely to be suitable hosts for specific pathogens. Ecological, phylogenetic, and life history traits are predictive of viral host status across mammalian species (Olival et al. 2017; Guy et al. 2020), as well as rates of cross-species viral transmission (Albery et al. 2020). Trait-based machine learning models are powerful tools for reservoir host prediction due to their ability to correct for sampling bias, perform out-of-sample prediction, and handle non-random missing data (Han et al. 2015). This approach can accurately predict host–virus interactions at various taxonomic scales: orthopoxviruses across all mammals (Tseng et al. 2025), ebolaviruses in African mammals (Sundaram et al. 2022), filoviruses across all bats (Han et al. 2016), and Nipah virus in a geographically relevant subset of bats (Plowright, Becker, Crowley, et al. 2019). The predictions generated by these models can then inform which putative host species are targeted for sampling in the field or, alternatively, in museum collections.

In this study, we present a framework for model-guided virus discovery in historical museum collections. We demonstrate this approach through two case studies focusing on viral RNA families with known bat reservoir hosts: coronaviruses (*Coronaviridae*) and paramyxoviruses (*Paramyxoviridae*). Both viral families are of growing human health concern. The *Coronaviridae* contains but is not limited to betacoronaviruses and alphacoronaviruses that cause various illnesses, including SARS, COVID-19, MERS, and several mild seasonal respiratory illnesses (Zhu et al. 2020). Paramyxoviruses include but are not limited to the zoonotic henipaviruses, including Hendra and Nipah virus, which cause fatal encephalitis in humans and domestic animals with no known treatments or human vaccines (Pomeroy et al. 2008). Both viral families occur across vertebrate taxa, but a wide diversity of coronaviruses and paramyxoviruses have been identified in bats, suggesting these volant mammals may be their ancestral hosts (Woo et al. 2006; Drexler et al. 2012). Consequently, bats have been heavily, though unevenly, sampled for these viruses, yielding enough viral data to train predictive models (Becker et al. 2019). However, both coronaviruses and paramyxoviruses have large spatial and taxonomic sampling gaps in bats (Cohen et al. 2023; Juman et al. 2025). Museum collections therefore offer promising opportunities to efficiently fill these gaps by targeting predicted host species found across the world and the bat phylogeny.

In our case studies of model-driven viral discovery, we leveraged the extensive and taxonomically diverse bat tissue collection housed at the Field Museum of Natural History (Chicago, USA). The depth and breadth of this collection also allowed us to test which sample types, preservation methods, and storage media are most conducive to RNA virus detection in museum tissue samples. First, we screened specimens for coronaviruses based on predictions from previously published models of betacoronavirus hosts across the whole order Chiroptera (Becker et al. 2022). For our paramyxovirus case study, we chose the best-performing approach—boosted regression trees (BRTs), a trait-based model—and applied it to a taxonomic subset of bats known to host zoonotic paramyxoviruses, the Old World fruit bats (family Pteropodidae) (Halpin et al. 2000; Chua et al. 2002). Taxonomically specific models outperform full models in predicting host–pathogen relationships (Dallas and Becker 2021), so we constrained our second case study to test this approach. More broadly, we sought to: 1) empirically test two predictive modeling approaches at different taxonomic scales; and 2) identify novel viruses and host–virus associations.

## Results

### Coronavirus host prediction and specimen screening

Our original betacoronavirus model predicted 412 bat host species that are not currently known to be betacoronavirus hosts (Table S1; Becker et al. 2022). We screened 68 of these species (806 samples) as well as an additional 524 samples of other bat species (including known hosts) based on the availability and quality of tissue samples at the FMNH. Overall, we screened 99 species of bats from 54 genera and nine families. This constituted 1330 samples collected between 1986 and 2023 in 10 countries across Africa, Asia, and South America. Our samples included heart (*n* = 76), liver (*n* = 117), kidney (*n* = 8), colon (*n* = 146), muscle (*n* = 25), bone fragment (*n* = 2), mixed tissue (*n* = 47), and fecal (*n* = 11) samples, in various preservation media: DMSO (*n* = 35), DNA/RNA Shield (*n* = 159), ethanol (*n* = 57), or flash-frozen without buffer (*n* = 113).

Of these 1330 specimens, 16 were positive for coronaviruses (Table 1). All RdRp sequences were recovered from colon samples (either flash-frozen or frozen in DNA/RNA Shield) collected from bats in Angola and the Democratic Republic of the Congo (DRC) between 2019 and 2023. RdRp sequences were recovered from two suspected hosts predicted by our model: *Rhinolophus fumigatus* (1/22 tested; 5% prevalence) and *Casinycteris argynnis* (1/7 tested; 14% prevalence). Samples from the other 66 predicted host species screened by PCR were negative. Nine *Hipposideros ruber* (a known host) yielded PCR-positive colons (9/16 tested; 56% prevalence), and we also detected coronaviruses in colons of one *Myonycteris torquata* (1/22 tested; 5% prevalence) and one *Rousettus aegyptiacus* (1/9 tested; 11% prevalence), two other previously known hosts. Two sequences were recovered from individuals not identified at the species level (*Hipposideros* spp. and *Neoromicia* spp.). The final sequence was recovered from one *Afropipistrellus grandidieri* (1/3 tested; 33% prevalence), a species that was not included in the original modeling study (as it was missing from the taxonomic backbone) but is nonetheless also a novel coronavirus host species. Other PCR products of the appropriate sequence lengths (i.e., ∼440 bp) were revealed by Sanger sequencing to be bat mRNA. All samples from species predicted as unlikely hosts (screened anyway because of available samples; *n* = 35) were negative.

**Table 1.** Positive samples from screening of museum specimens for coronaviruses and paramyxoviruses. Further details on locality can be found in Supplementary Data S1. All positive samples were colon samples.

| Catalog number | Taxon | Country | Generalized locality | Year | Model Prediction | Viral family | Buffer |
| --- | --- | --- | --- | --- | --- | --- | --- |
| FMNH 238753 | *Rhinolophus fumigatus* | Angola | Bié Province | 2019 | Predicted host | CoV | None (flash-frozen) |
| FMNH 238754 | *Hipposideros ruber* | Angola | Bié Province | 2019 | Known host | CoV | None (flash-frozen) |
| FMNH 244350 | *Hipposideros ruber* | Angola | Namibe Desert | 2023 | Known host | CoV | DNA/RNA Shield |
| FMNH 244367 | *Hipposideros ruber* | Angola | Namibe Desert | 2023 | Known host | CoV | DNA/RNA Shield |
| FMNH 244593 | *Hipposideros ruber* | DRC | Tshopo | 2023 | Known host | CoV | DNA/RNA Shield |
| FMNH 244335 | *Hipposideros ruber* | Angola | Namibe Desert | 2023 | Known host | CoV | DNA/RNA Shield |
| FMNH 244336 | *Hipposideros ruber* | Angola | Namibe Desert | 2023 | Known host | CoV | DNA/RNA Shield |
| FMNH 244340 | *Hipposideros* spp. | Angola | Namibe Desert | 2023 | Not in model | CoV | DNA/RNA Shield |
| FMNH 244341 | *Hipposideros ruber* | Angola | Namibe Desert | 2023 | Known host | CoV | DNA/RNA Shield |
| FMNH 244343 | *Hipposideros ruber* | Angola | Namibe Desert | 2023 | Known host | CoV | DNA/RNA Shield |
| FMNH 244580 | *Myonycteris torquata* | DRC | Tshopo | 2023 | Known host | CoV | DNA/RNA Shield |
| FMNH 244584 | *Rousettus aegyptiacus* | DRC | Tshopo | 2023 | Known host | CoV | DNA/RNA Shield |
| FMNH 242739 | *Afropipistrellus grandidieri* | Angola | Serra da Neve | 2022 | Not in model | CoV | DNA/RNA Shield |
| FMNH 244609 | *Neoromicia* spp. | DRC | Tshopo | 2023 | Not in model | CoV | DNA/RNA Shield |
| FMNH 244594 | *Hipposideros ruber* | DRC | Tshopo | 2023 | Known host | CoV | DNA/RNA Shield |
| FMNH 244562 | *Casinycteris argynnis* | DRC | Tshopo | 2023 | Predicted host | CoV | DNA/RNA Shield |
| FMNH 244565 | *Epomops franqueti* | DRC | Tshopo | 2023 | Predicted host | PmV | DNA/RNA Shield |
| FMNH 244607 | *Glauconycteris humeralis* | DRC | Tshopo | 2023 | Not in model | PmV | DNA/RNA Shield |
| FMNH 238751 | *Rhinolophus fumigatus* | Angola | Bié Province | 2019 | Not in model | PmV | None (flash-frozen) |

Based on the top BLAST hits to sequences in GenBank, ten RdRp sequences were most closely related (87.72%–96.74%, mean 89.55%) to those in the *Alphacoronavirus* genus, and six were most closely related (95.01%–100.00%, mean 98.64%) to sequences in the *Betacoronavirus* genus (Table 2). Eight of the sequences (50%) had <90% similarity to known sequences in GenBank and are thus novel bat coronaviruses (Anthony et al. 2017). Of the 16 RdRp sequences, we were able to align the 13 most complete sequences and generate a phylogeny (Fig. 1). The *Rhinolophus* sequence (FMNH 238753) clusters with sarbecovirus sequences closely related to SARS-CoV-2.

**Table 2.**
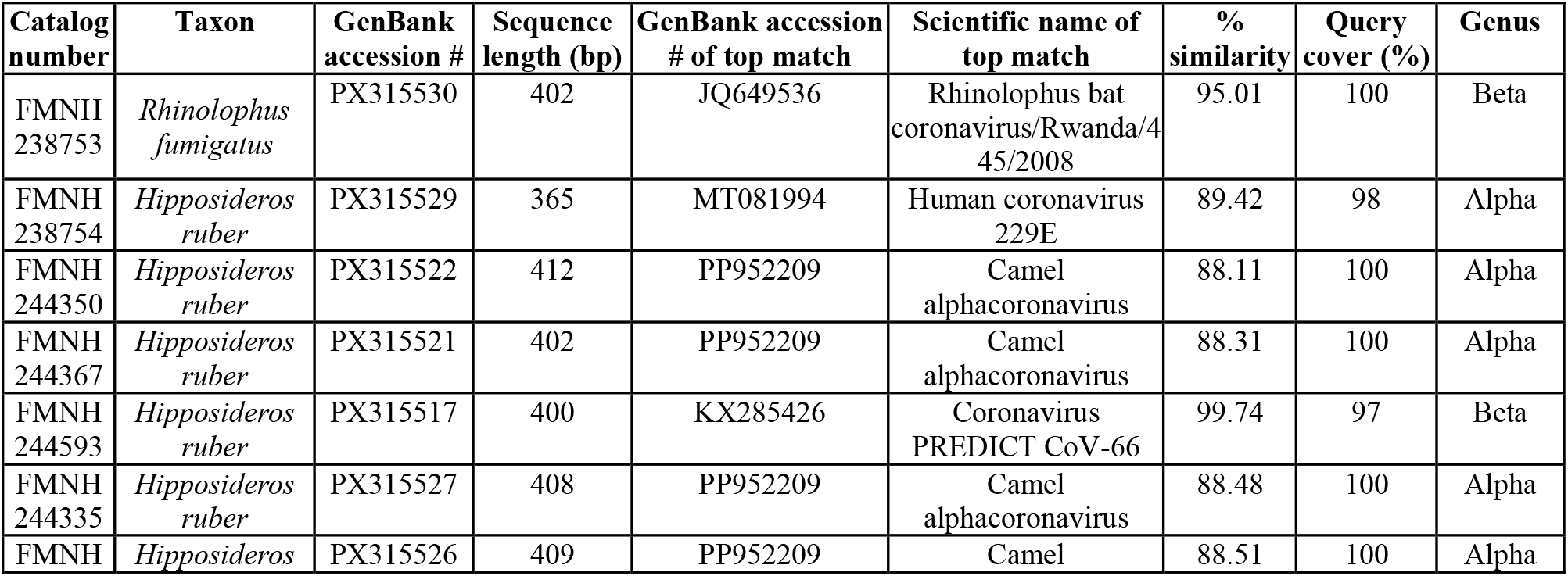
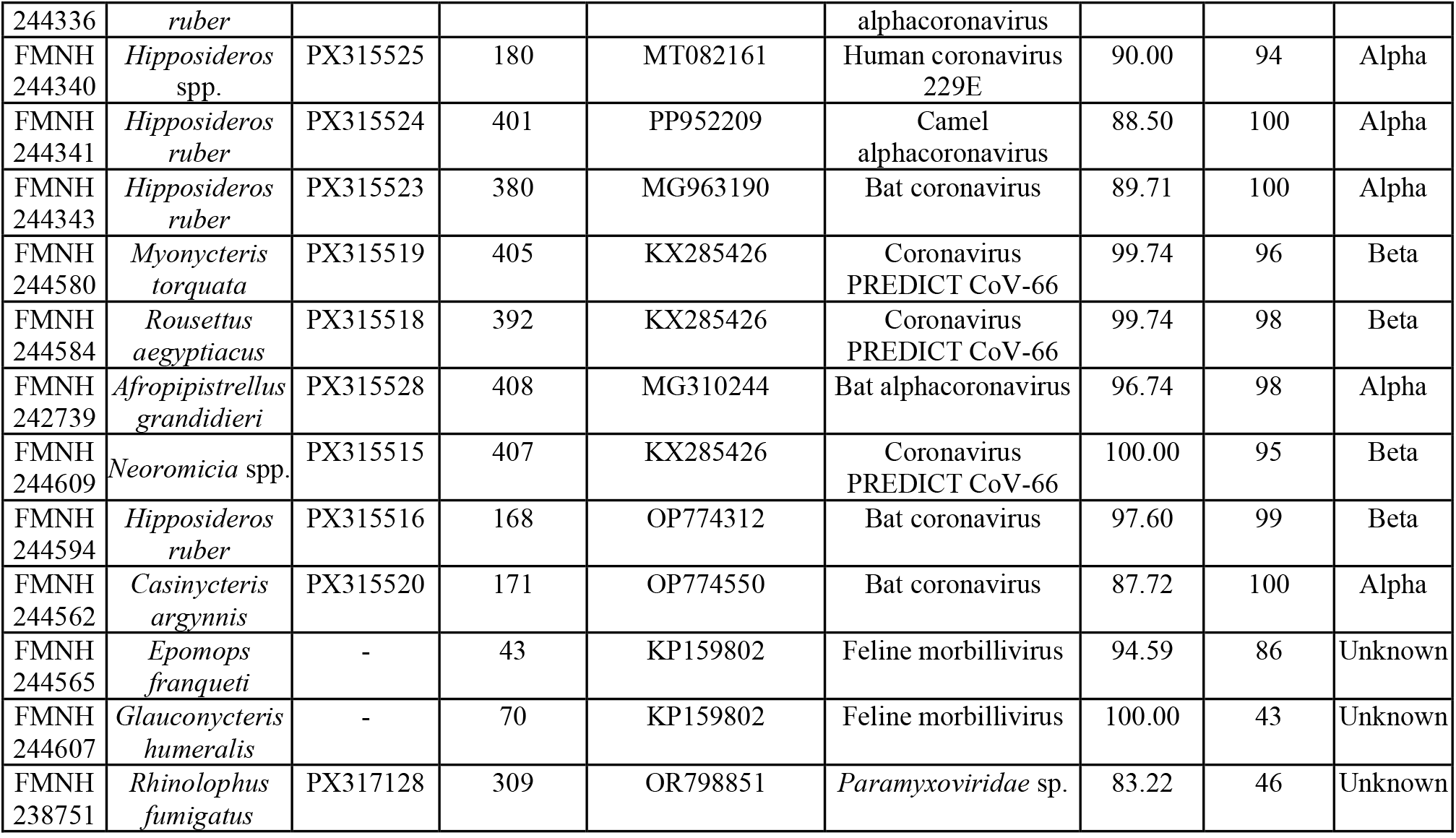
Sequences obtained from museum specimens and BLAST results from July 9 2025. When top values are identical, we report the highest bat match. Matches were sorted first by E value and then by percent identification.

**Fig. 1.**
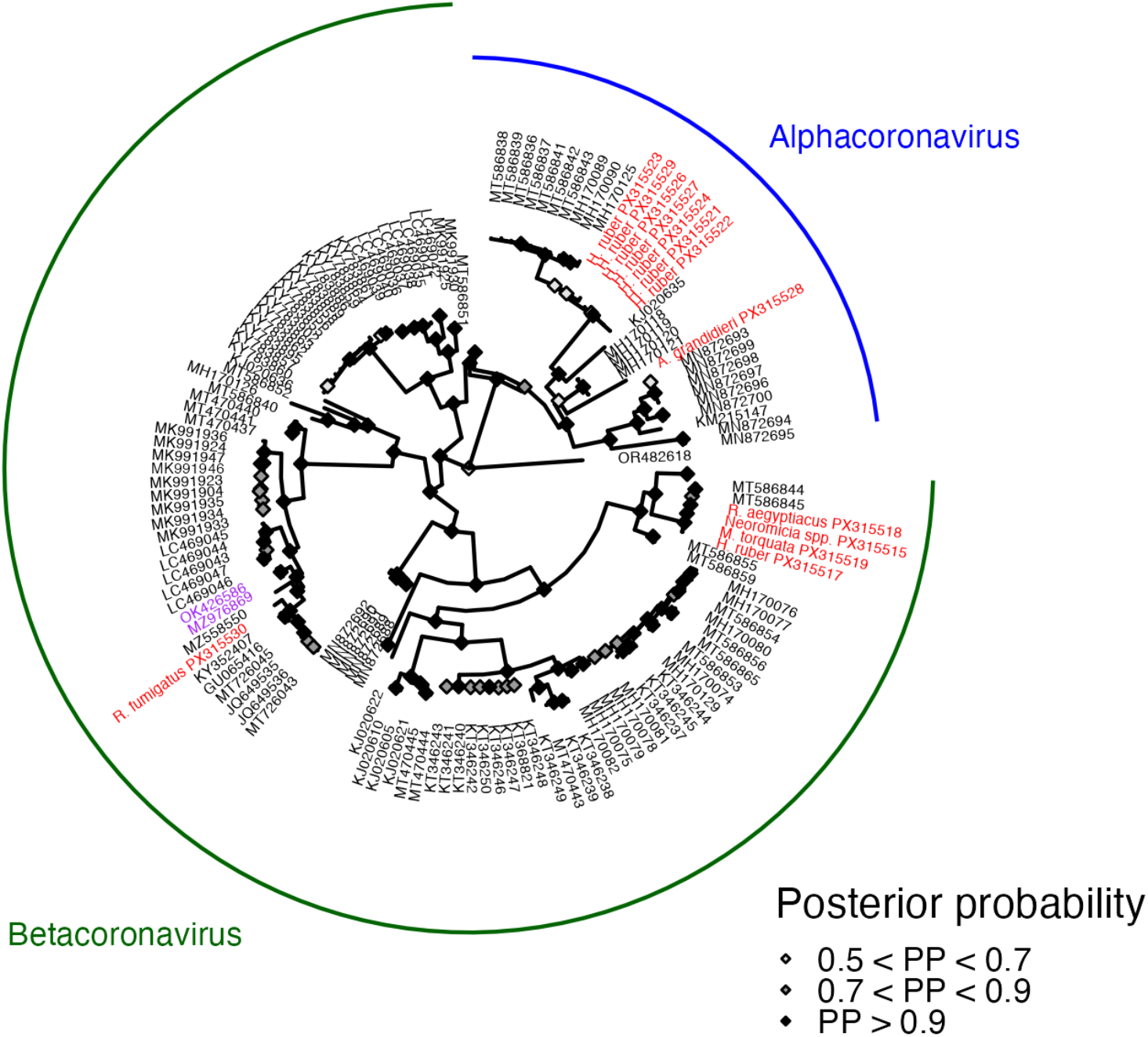
Circular phylogeny for the paramyxovirus L gene estimated using MrBayes with 1,000,000 iterations, a GTR+I+G model, and 20% burn-in (Ronquist et al. 2012). Recovered sequences are labeled in red, while SARS-CoV-2 isolates are labeled in purple.

### Paramyxovirus host prediction and specimen screening

Our BRT analysis identified paramyxovirus PCR-positive pteropodid bat species with up to 92% accuracy (AUC = 0.92, SE = 0.01; corrected AUC = 0.79, SE = 0.02). Model outputs and hyperparameters are listed in Table S3. The top six variables identified by the model as influential predictors of host status were virus-related PubMed citations, total range area, all PubMed citations, adult body length, ear length, and evolutionary distance (equal splits); all other predictors had <3% importance (Table S3). Partial dependence plots revealed that both citation counts, as well as range area and ear length, were positively associated with paramyxovirus host status, while evolutionary distance was negatively associated with host status. Adult body length was more non-linearly associated with host status, suggesting that pteropodid species of small-to-medium sizes are more likely to be hosts (Fig. 2). Based on this trait profile, the most likely “novel” host species (with no previous PCR paramyxovirus detections) are *Cynopterus sphinx, Epomops franqueti, Cynopterus brachyotis, Epomophorus wahlbergi, Rousettus madagascariensis*, and *Macroglossus minimus*, all of which had positivity probabilities above 20% (Table S4). To account for the fact that species with higher citations are more likely to be predicted as hosts (Fig. 2A,C), we also reran our predictions when fixing total and virus-related citations at their mean. Here, the three “novel” predicted species were *Epomops franqueti, Epomophorus wahlbergi*, and *Macroglossus minimus*, all of which still had positivity probabilities above 15% (Table S4).

**Fig. 2.**
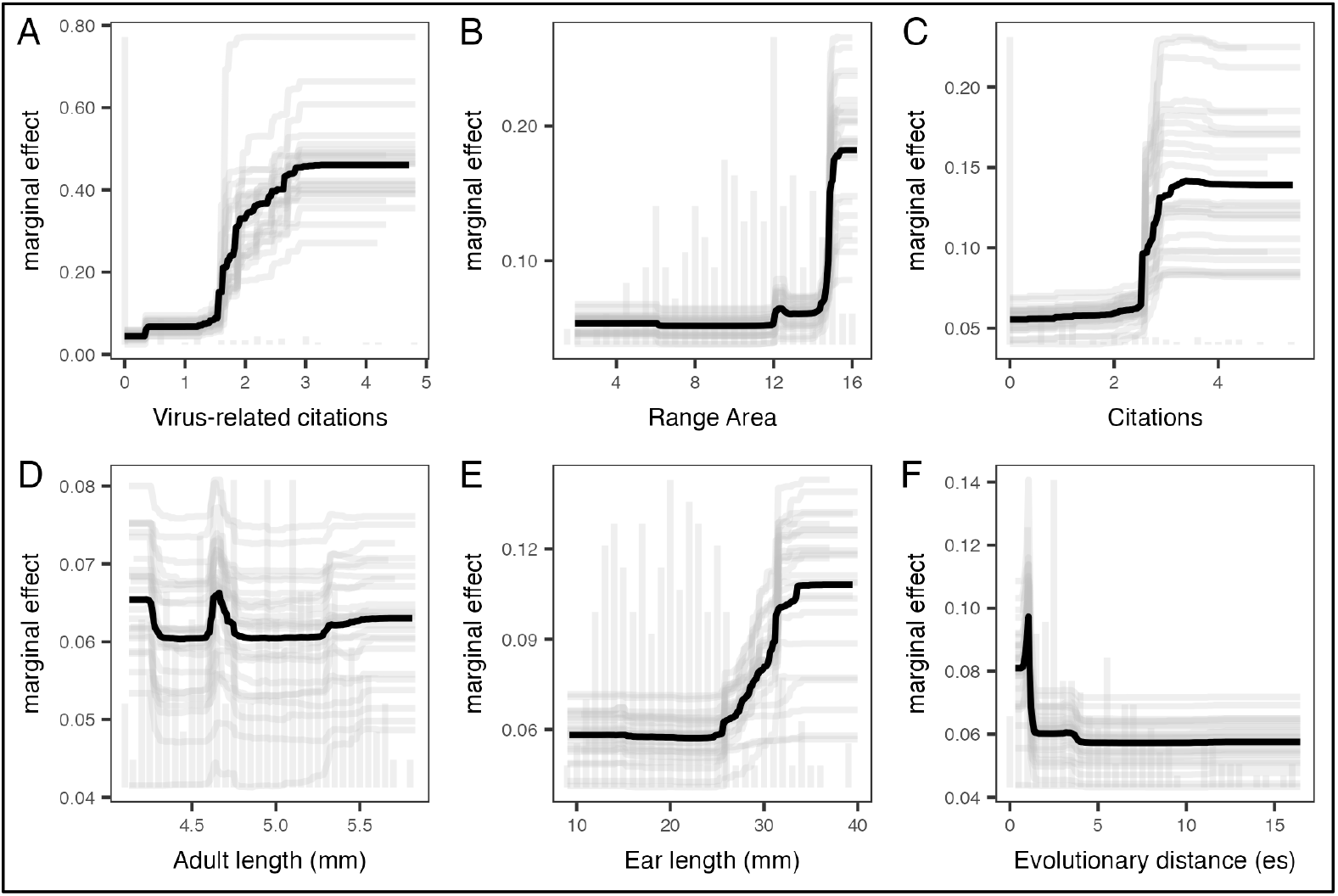
Partial dependence plots for the six most influential predictors of paramyxovirus PCR positivity in Pteropodidae: A) natural log of virus-related PubMed citations; B) natural log of total range area (km^2^); C) natural log of all PubMed citations; D) natural log of adult body length (mm); E) ear length (mm); F) evolutionary distance (equal splits).

We screened these three suspected host species (*Epomops franqueti, n* = 20; *Epomophorus wahlbergi, n* = 27; *Macroglossus minimus, n* = 89), as well as an additional 355 specimens of other bat species (including known hosts) based on the availability and quality of tissue samples at the FMNH. Overall, we screened 36 species belonging to 27 genera and six families. Given our earlier success screening colon samples in DNA/RNA Shield for coronaviruses, we prioritized similar samples for further paramyxovirus screening. In total, we screened 491 samples collected between 1989 and 2023 in Angola, the DRC, Tanzania, and the Philippines. Our samples included colon (*n* = 221), spleen (*n* = 40), and mixed tissue samples (*n* = 230), preserved either in DNA/RNA Shield (*n* = 129) or flash-frozen without buffer (*n* = 230).

Of these 491 samples, three were positive for paramyxoviruses (Table 1). All three L gene sequences were recovered from colon samples (flash-frozen or frozen in DNA/RNA Shield) collected from bats in Angola and DRC between 2019 and 2023. One sequence was recovered from *Epomops franqueti*, one of the three suspected host species predicted by our model, after correcting for citation counts. We also recovered two sequences from *Rhinolophus fumigatus* and *Glauconycteris humeralis*, insectivorous species that were not included in our pteropodid BRT but are also novel paramyxovirus hosts (these samples had been previously screened for coronaviruses and the extracted RNA was reused for paramyxovirus screening). Other PCR products of the appropriate sequence lengths (200-500 bp) were revealed by Sanger sequencing to be bat mRNA. All samples from species predicted as unlikely hosts (screened anyway because of available samples; *n* = 35) were negative.

The two shorter, partial sequences (from *E. franqueti* and *G. humeralis*) could not be fully aligned and therefore were excluded from our phylogenetic analysis. However, BLAST revealed that these sequences most closely matched to a feline morbillivirus (Table 2). The *R. fumigatus* sequence (FMNH 238751) most closely matched a distantly related (<90% similarity) paramyxovirus sequence isolated from a Chinese *Rhinolophus* kidney (Table 2), suggesting that this sequence is from a novel bat virus. We were able to align this latter sequence to reference sequences (Zhu et al. 2022) and generate a phylogeny based on the paramyxovirus L gene (Fig. 3). Our *R. fumigatus* sequence clusters with other unclassified bat paramyxoviruses in the *Orthoparamyxovirinae* subfamily (Fig. 3).

**Fig. 3.**
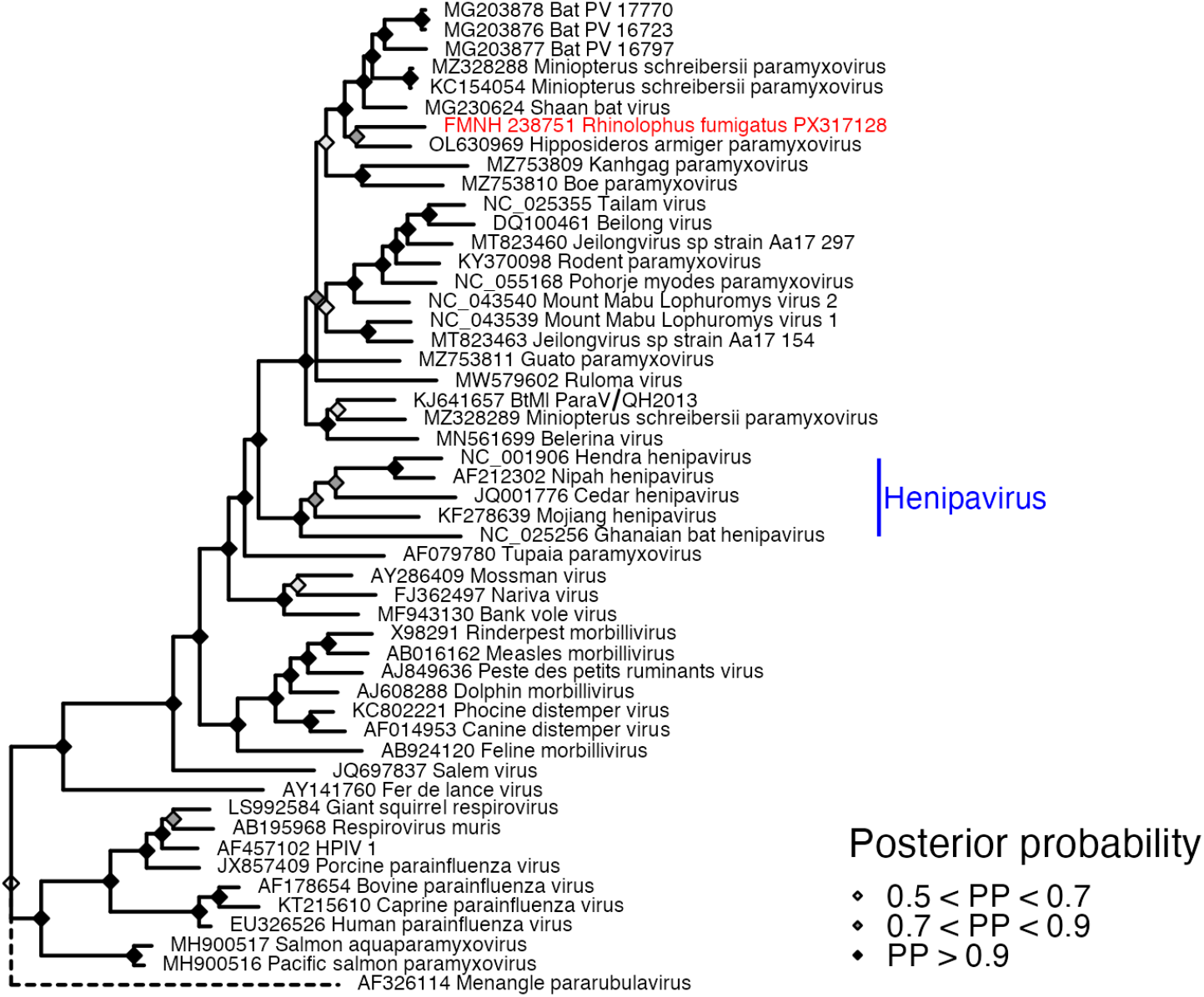
Phylogeny for the paramyxovirus L gene estimated using MrBayes with 1,000,000 iterations, a GTR+I+G model, and 20% burn-in (Ronquist et al. 2012). Comparative sequences were derived from Zhu et al. (2022). The recovered sequence from FMNH 238751 (*Rhinolophus fumigatus*) is labeled in red, with the henipavirus clade labeled in blue.

## Discussion

Leveraging museum collections for pathogen discovery is a topic of growing interest (Thompson et al. 2021; Colella et al. 2021; Paul et al. 2025). But attempts to screen museum specimens for pathogens remain rare, as such research is restricted by practical and logistical challenges, including the sheer depth and taxonomic diversity of collections (Johnson et al. 2023). Combining predictive analytics with these collections offers an efficient and productive method for selecting specimens. This focused approach has the potential to raise detection rates relative to random screening, enabling the characterization of new pathogens and further understanding of the host ranges of known pathogens.

We conducted two case studies to evaluate the utility of museum specimens for empirically testing model-generated predictions of RNA virus host suitability. Sixteen coronavirus sequences (eight of which were novel with <90% similarity to known sequences) were recovered from museum specimens collected from 2019 to 2023, including from three novel coronavirus hosts: *Rhinolophus fumigatus, Casinycteris argynnis*, and *Afropipistrellus grandidieri*. Both *R. fumigatus* and *C. argynnis* were suspected betacoronavirus hosts in our predictive model (Becker et al. 2022; Table S1), while *A. grandidieri* was not included in the model but was tested because of the available tissue sample. Based on preliminary phylogenetic analyses and GenBank matches, we recovered a betacoronavirus and an alphacoronavirus sequence from *R. fumigatus* and *A. grandidieri*, respectively, with a partial alphacoronavirus sequence recovered from *C. argynnis* (Table 2). Of particular interest for pandemic risk assessment, the *R. fumigatus* betacoronavirus RdRp sequence was closely related to that from sarbecoviruses, including SARS-CoV and SARS-CoV-2 (Table 2; Fig. 1). This specimen was collected from the Bié Province in Angola in June 2019, from a site near agro-pastoral land and human communities (Supplementary Data S1).

This model-guided viral discovery approach was also successful for paramyxoviruses, as we recovered sequences from a novel paramyxovirus host predicted by our pteropodid-focused model (*Epomops franqueti*) as well as two novel hosts not included in our model (*R. fumigatus* and *Glauconycteris humeralis*). All positive specimens were collected from sub-Saharan Africa, where there have been no confirmed paramyxovirus spillover events and bats remain undersampled for these viruses (Juman et al. 2025). However, serological evidence from Cameroon hints at potential henipavirus exposure in humans (Pernet et al. 2014). The paramyxovirus-positive *R. fumigatus* was collected from the same site as the coronavirus-positive *R. fumigatus* (Supplementary Data S1). The *E. franqueti* specimen was also collected near a main road, and this species’ occurrence is generally associated with human activity (Olivero et al. 2020), suggesting opportunities for spillover. Phylogenetic analysis based on the L gene of the most complete sequence (*R. fumigatus*) suggested that it falls within the *Orthoparamyxovirinae* subfamily, which also includes the zoonotic Hendra and Nipah henipaviruses (Table 2; Fig. 3). It is most closely related to viruses in the *Jeilongvirus* genus, some of which have a broad range of susceptible host species and therefore zoonotic potential (DeRuyter 2024). However, this sequence had <90% similarity to known sequences, suggesting that it belongs to a novel bat paramyxovirus (Anthony et al. 2017). Complete genomic sequencing of these PCR-positive museum samples will shed further light on their phylogenetic relatedness to known pathogens of concern, which will be critical for understanding the evolution and transmission potential of these novel paramyxoviruses in light of the high case fatality rates and frequent spillover of Asian and Australian paramyxoviruses (Pomeroy et al. 2008; Juman et al. 2025).

Our research validates the approach of trait-based predictive modeling for generating ranked lists of suspected hosts; in both case studies, at least one suspected novel host was confirmed through empirical testing, and none of the samples from species predicted as unlikely hosts (*n* = 35 for each case study) were PCR-positive. New host–virus associations can be used to inform and iteratively refine the original model (Becker et al. 2022). While many studies have modeled suspected hosts in other mammal–virus systems (e.g., Han et al. 2016; Plowright, Becker, Crowley, et al. 2019; Sundaram et al. 2022; Fischhoff et al. 2021), few have gone on to empirically test these predictions, and those that have done so have employed field surveys (Gokhale et al. 2021) or experimental studies (Seifert et al. 2020; Lasso et al. 2025). Here, we present the first case studies of validating host–virus predictions with natural history museum collections. Two of 68 screened suspected coronavirus hosts were positive by PCR and one of three screened suspected paramyxovirus hosts were positive by PCR. While the majority of our suspected host species were negative when screened in collections, this may be a function of sample size, preservation method, and low viral prevalence. These predicted hosts should remain potential targets for future field- or museum-based sampling.

Our findings also underscore the value of museum collections for viral discovery and surveillance more generally. The use of natural history collections for viral surveillance and host identification has gained more attention in recent years (Colella et al. 2021), yet these collections remain largely underutilized resources for viral discovery, particularly for RNA viruses. Our study is the first to establish that RNA from coronaviruses and paramyxoviruses, two viral families that cause high disease burdens in humans, can be recovered from museum tissues, confirming the utility of these collections in discovering new hosts. Host–virus associations in museum collections can also provide data about the geographic ranges and tissue tropism of viruses, as well as the timing of their emergence into new species. Critically, they also offer a cost-effective and tractable avenue for addressing geographic and taxonomic sampling gaps, potentially offering increased safety in more controlled settings compared to the logistical and biosafety challenges of traditional field-based approaches (Thompson et al. 2021). Here, we used museum collections to screen for coronaviruses and paramyxoviruses in bats from sub-Saharan Africa, which have been overlooked relative to Chinese bats and South Asian and Australian bats, taxa that have been disproportionately screened for coronaviruses and paramyxoviruses, respectively (Cohen et al. 2023; Juman et al. 2025).

Museums also serve as time machines for studying disease emergence. Our discovery of a SARS-like coronavirus in an Angolan bat from June 2019 suggests that betacoronaviruses with pandemic potential have been circulating in regions with potential bat–human contact since before the SARS-CoV-2 pandemic. None of the positive bats were collected from primary and protected forests. While there have been no known betacoronavirus or paramyxovirus spillovers in sub-Saharan Africa yet, risk modeling suggests that West and Central Africa are possible hotspots for future zoonotic emergence events (Forero-Muñoz et al. 2024). Our findings reinforce that sub-Saharan Afrotropical bats and their viruses warrant more research focus in future viral discovery studies. Nearby human communities could be considered targets for increased disease surveillance and capacity building (Worsley-Tonks et al. 2022). This includes investment in establishing and maintaining museum collections, biorepositories, and laboratories in the Global South to enable proactive and efficient in-country screening of wildlife for pathogens (Colella et al. 2020).

However, museum collections also pose unique challenges for viral surveillance studies. Recovery of viral sequences is highly dependent on preservation method, storage medium, and time elapsed since collection. All positive samples in this study were stored in a cryogenic tissue storage facility, either flash-frozen immediately after collection or preserved in DNA/RNA Shield. The oldest specimen screened in our study was collected in 1986, but our earliest positive results came from bats collected in 2019. This is likely a function of the fact that collectors have only recently begun flash-freezing or using appropriate buffers for downstream viral diagnostics. Additionally, the rapid degradation of RNA creates a further challenge; specimens must be immediately sampled upon collection for RNA recovery to later be feasible. Whenever possible, museums and collectors should consider adopting collection and storage practices that produce samples amenable to longer-term RNA preservation and future viral research (Thompson et al. 2021). However, recent methodological advances in obtaining DNA and RNA out of ancient and formalin-fixed specimens could unlock a broader temporal range of museum specimens for future viral screening (Speer et al. 2022; Hahn et al. 2024; Hämmerle et al. 2024).

In summary, our study provides a framework for—and shows the promise of—model-guided viral discovery in natural history museum collections (Fig. 4). This approach can be adapted for future studies of other pathogens and customized to the taxonomic or geographic scales of interest (e.g., all bats for betacoronaviruses, only pteropodid bats for all paramyxoviruses). Once any novel host–pathogen associations are identified, further ecological and experimental studies of transmission dynamics are required to determine whether these hosts are true reservoirs or only incidental hosts (Viana et al. 2014). Such data can then feed back into improved modeling efforts and stronger inference about the role of such species for multi-host pathogens (Becker et al. 2020). Museum collections are limited in that they provide a snapshot of a host–pathogen association at one point in space and time; still, they offer a valuable starting point for investigation. Further collaboration between natural history museums, modelers, and disease ecologists is critical for facilitating this line of work (Paul et al. 2025), as is continued and mindful collection, preservation, and digitization. Specimens are an investment in future science and a unique record of global change, potentially enabling us to study the responses of hosts and microbes to unprecedented anthropogenic impacts.

**Fig. 4.**
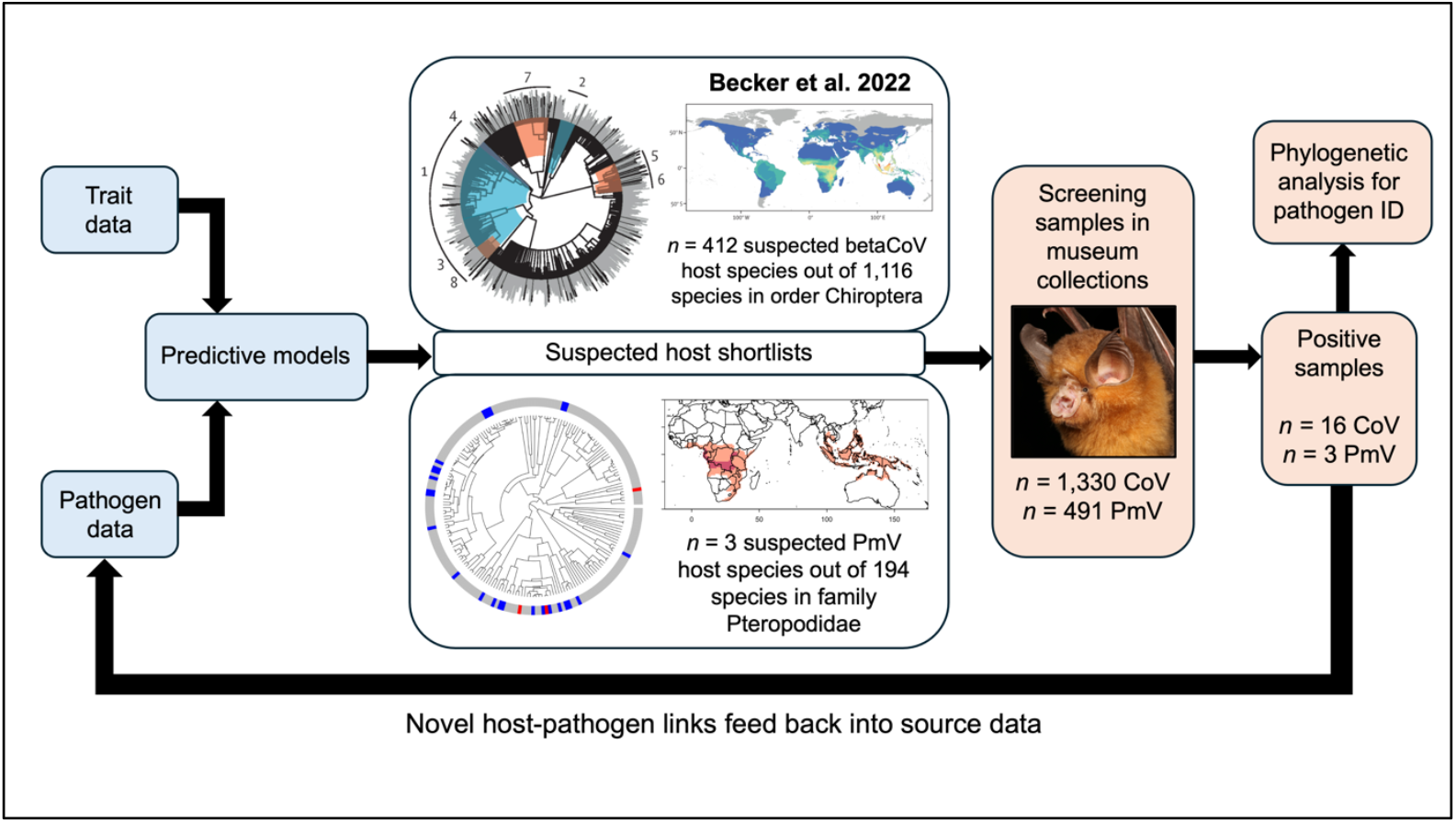
Flowchart outlining the process of model-guided coronavirus and paramyxovirus discovery in natural history museum collections. The suspected host shortlists in this study were generated by separate machine learning models (Becker et al. 2022 for the coronavirus predictions). The phylogenetic positions (in red) and species richness of predicted host species inform which museum collections are targeted for screening.

## Methods

### Predictive modeling

For betacoronavirus host prediction, we used the results of a previously published study (Becker et al. 2022), which consolidated eight models (four network-based, three trait-based, and one hybrid) to generate predictions about likely betacoronavirus hosts across all bats (Table S1). When these models were validated with updated betacoronavirus surveillance data during the COVID-19 pandemic, trait-based models consistently outperformed network-based approaches. Therefore, for subsequent paramyxovirus host prediction, we chose one of the best-performing trait-based models in the first case study— boosted regression trees (Elith et al. 2008)—and constrained it to the bat family Pteropodidae, following an earlier study of taxonomically specific models (Dallas and Becker 2021).

We first assembled data on Pteropodidae–paramyxovirus associations from the literature and created the pteroparamyxo database (Juman et al. 2025) for this and future modeling studies. We then included any additional associations from the VIRION database (Carlson et al. 2022; accessed on 3 August 2023). In total, nine different types of associations were recorded as binary variables: molecular (PCR) evidence of association, serological evidence of association, or successful viral isolation, for three different taxonomic groups (*Paramyxoviridae* overall and subfamilies *Orthoparamyxovirinae* and *Rubulavirinae*). In the final model, we used PCR detections and viral isolation at the highest taxonomic level (*Paramyxoviridae*) to make broad host predictions across the entire viral family. We excluded serological detections, which may reflect antigen cross-reactivity and are therefore less specific (Gilbert et al. 2013).

We then compiled a suite of ecological, morphological, life history, and geographic traits from the COMBINE mammalian trait database (Soria et al. 2021) and IUCN (2025), supplemented by other published databases of bat-specific functional and morphological traits (Giannini 2019; Guy et al. 2020; Cosentino et al. 2023). We used only reported traits rather than imputed data. We also added two measures of evolutionary distance (“fair proportion” and “equal splits”) calculated using a phylogeny of the Pteropodidae (Upham et al. 2019) as well as dummy variables for each of 43 bat genera. Finally, as a proxy for study effort, we included two citation count metrics extracted from PubMed using the *easyPubMed* R package (Fantini 2019) on 22 February 2024 (all citations per bat species and all virus-related citations per bat species). We removed any variables that were either completely homogenous across all bat species and/or missing data for greater than 40% of the species (Fig. S1). Our final dataset included 83 predictors for 194 pteropodid bat species harmonized to a taxonomic backbone (Upham et al. 2019). All included predictors, along with their descriptions, coverage, and citations, are listed in Table S2.

Of the 194 pteropodid bat species, 25 had published PCR associations with paramyxoviruses (coded as 1 in our binary response variable; all unsampled or negative bats were coded as 0), and we trained a BRT model via the *gbm* package (Greenwell et al. 2022) to classify these known positives from unsampled or negative species. This is a pattern recognition, hypothesis-generating approach that identifies predictors associated with a binary response variable (i.e., paramyxovirus PCR positivity) and extrapolates those patterns to negative or unsampled species to predict their likelihood of viral positivity. We trained our BRTs on 80% of the complete dataset, using 25 splits, 3000 trees with a Bernoulli error distribution, and five-fold cross-validation to prevent overfitting (Elith et al. 2008). We calculated the corrected area under the curve (AUC) as a performance metric on both our training and test dataset using target shuffling methods (Han et al. 2016). Model parameters were selected based on comparisons of the test AUC, sensitivity, and specificity of models across a grid search of factorial combinations of hyperparameters (Fig. S2). Final hyperparameter values and model outputs are reported in Table S3. Each variable was assigned a score indicating its relative contribution to classification, with higher values indicating stronger influence on the model and the sum of all relative influence scores adding up to 100. The contribution of each variable can be visualized using a partial dependence plot, which indicates the effect of the predictor on the response when all other predictors are averaged. We calculated the final relative influence and partial dependence of each variable by averaging these outputs across all 25 model splits. Finally, we predicted the probability of paramyxovirus hosting across unsampled or negative bat species, including predictions where both citation count measures were held at their mean (Becker et al. 2022; Mull et al. 2022; Tseng et al. 2025). We likewise averaged these predictions for each species across the 25 model splits and, for consistency with the betacoronavirus model (Becker et al. 2022), we used a 90% sensitivity threshold with the *PresenceAbsence* package to stratify results into binary predictions of suspect and non-suspect paramyxovirus hosts (Freeman and Moisen 2008). This package can be applied to the raw probability values from our model to examine other potential cutoffs (Table S4).

### Specimen selection and viral screening

We empirically tested our model predictions by screening frozen bat tissue samples from specimens housed at the Field Museum of Natural History (FMNH), Chicago, USA. Samples were selected based on the results of our models (Table S1; Table S4), with species predicted to be suitable betacoronavirus or paramyxovirus hosts (following a 95% sensitivity cutoff for predicted probabilities) prioritized for screening. Sample selection was also influenced by the availability of tissues in appropriate preservatives and storage media, with flash-frozen samples or samples preserved in appropriate buffer (e.g., DNA/RNA Shield) prioritized for having the highest likelihood of RNA virus detection. Based on preliminary evidence of tissue tropism, we focused on colon samples for coronaviruses (Mols et al. 2023) and spleen and colon samples for paramyxoviruses (Juman et al. 2025). High-quality tissue samples were screened for both coronaviruses and paramyxoviruses to maximize reagents and previously extracted RNA or converted cDNA, even if outside the taxonomic scope of the case studies (e.g., some insectivorous bats were screened for paramyxoviruses, as RNA had already been extracted for the betacoronavirus case study). Ultimately, 1330 samples were screened for coronaviruses, and 491 samples were screened for paramyxoviruses.

Tissues were subsampled from the FMNH cryogenic collection. For high-yield tissues (spleen and liver), 2 mg of tissue was removed and immediately stored in 200 μL DNA/RNA Shield (for every other tissue type, 5 mg of tissue was subsampled). Subsequent molecular analyses were conducted at the FMNH Pritzker Laboratory for Molecular Systematics and Evolution. Total nucleic acids were extracted from tissues using the Qiagen AllPrep DNA/RNA 96 Kit following the manufacturer’s protocol. An extra DNase treatment was used, following the Qiagen RNeasy protocol. Tissues from recent collection trips, stored in Zymo DNA/RNA Shield, were instead extracted using the Zymo Quick-DNA/RNA MagBead Kit with internal DNase treatment.

RT-PCR was performed using the SuperScript III One-Step RT-PCR System with Platinum Taq polymerase to target a 440 bp region of the betacoronavirus RdRp gene and a 200-500 bp region of the paramyxovirus L gene. For the coronavirus case study, RT-PCR followed published protocol (Waruhiu et al. 2017) using one-fourth reactions and 4 μL of RNA template. The thermal profile for coronavirus RT-PCR was 55°C for 30 min; 94°C for 2 min; 30 cycles of 94°C for 15 s, 48.4°C for 30 s, 68°C for 60 s; and a final extension at 68°C for 5 min. The second coronavirus PCR was performed using Platinum Taq Enzyme (Invitrogen) with 1 µL of cDNA following Waruhiu et al. (2017) and using 25 μL reactions. The thermal profile was as follows: 94°C for 5 min; 35 cycles of 94°C for 15 s, 51.8°C for 30 s, 72°C for 30 s; and a final extension at 72°C for 10 min. For the paramyxovirus case study, RT-PCR followed published protocol (Tong et al. 2008) using one-fourth reactions and 2.5 μL of RNA under the following thermal conditions: 60°C for 1 min; 47°C for 30 min; 94°C for 2 min; 40 cycles of 94°C for 15 s, 50°C for 30 s, 72°C for 30 s; and a final extension at 72°C for 7 min. The second paramyxovirus PCR was performed using Platinum Taq Enzyme (Invitrogen) with 1 µL of cDNA following Tong et al. (2008) and using 25 μL reactions. The thermal profile was as follows: 94°C for 2 min; 40 cycles of 94°C for 15 s, 50°C for 30 s, 72°C for 30 s; and a final extension at 72°C for 7 min. All PCRs were run on a Bio-Rad CFX96 Touch Real-Time Detection System.

PCR products were visualized with electrophoresis in a 2% low-melt agarose gel and bands at the expected length were extracted using the QIAquick gel extraction kit (Qiagen). Next, 100 ng of PCR product was dehydrated for a 5 μL cycle sequencing reaction using 2 μL of BigDye Terminator 3.1 Ready Reaction Mix (ABI) and 3 μL of 5 μM primer. The cycle sequencing thermal profile followed the manufacturer’s instructions. Samples were purified using standard ethanol/EDTA precipitation and resuspended in 10 uL of HiDi formamide. Samples were then sequenced on an ABI 3130 Genetic Analyzer.

### Phylogenetic analyses

After obtaining sequences, we reviewed chromatograms and trimmed primers in Geneious Prime (Kearse et al. 2012). When possible based on sequence quality, we generated consensus sequences using both forward and reverse reads. We then used NCBI BLAST to assess similarity with existing sequences on GenBank. Sequences with >90% identification with existing sequences in GenBank were considered members of the same clade (Anthony et al. 2017). We aligned our sequences with RdRp and L gene sequences from previously published genomes (Zhu et al. 2022) using Clustal Omega in Geneious and NGPhylogeny.fr (Lemoine et al. 2019); final alignments are provided as supplementary files. Phylogenetic trees were estimated using MrBayes 3.2.6, with a GTR+I+G model, 20% burn-in, and 1,000,000 iterations (Ronquist et al. 2012). A gammacoronavirus sequence and pararubulavirus sequence were used as outgroups for the coronavirus and orthoparamyxovirus phylogenies, respectively.

## Data and Code Availability

All specimens screened in this study are listed in Supplementary Data S1, also archived on Zenodo (10.5281/zenodo.17094222). Coronavirus RdRp sequences are available on GenBank under accessions PX315515–30 and the paramyxovirus L gene sequence from FMNH 238751 is available on GenBank under accession PX317128. Owing to the small size (<100 bp) of two of the paramyxovirus L gene sequences (FMNH 244607, FMNH 244565), we provide these sequence data in the supplement and on Zenodo. Nucleotide alignments for phylogenetic trees are also included in the supplement and on Zenodo. Data for pteropodid model training can be found in the pteroparamyxo database (https://github.com/mayajuman/pteroparamyxo). All code for pteropodid model development is housed in a GitHub repository (https://github.com/mayajuman/pteroparamyxo-BRT).

## Acknowledgments

We thank the curators and collection staff at the Field Museum of Natural History for their assistance with this project, including Lauren Johnson, Lawrence Heaney, Bill Stanley, and Anderson Feijó. We also thank Hendrik Poinar for consultation on laboratory methods. MMJ was supported by a Verena Fellow-in-Residence Award funded by NSF DBI 2515340 and a Gates Cambridge Scholarship enabled by grant OPP1144 from the Bill & Melinda Gates Foundation. MMM was supported by NSF CAREER Grant 2238801. Further funding was provided by the James L. Patton Award from the American Society of Mammalogists, a Field Museum Science Innovation Award, and a Negaunee Seed Fund Grant. This project was also supported by funding from the Cambridge Centre for Data-Driven Discovery and Accelerate Programme for Scientific Discovery, made possible by a donation from Schmidt Futures. Collecting and exporting permits in Angola were provided by the Instituto Nacional da Biodiversidade e Áreas de Conservação (74/INABC.MINAMB/2019, 12/INBAC.MINAMB/2022, 72INBC.MINAMB/2023, 62INBC/MINAMB/2023, 05INBC/MINAMB/2023) and in the DRC by the Universite de Kisangani and the Ministre Provincial des Forets, Environment, Agriculture, Peche (N/Ref FS/VDR/NBS/236/2023). Fieldwork in Serra da Neve (Angola) was supported by National Geographic Society Explorer Grant (NGS-73084R-20) to LMPC. Important field assistance was provided by Suzana Bandeira, Mariana Marques, Diogo Parrinha, Varito Baptista, and Ben D. Marks (Angola) and Jacques Tanzito, Glodi Lotana, Robert Abane, Charles Balikage, Fidiele Mushagalusa, Gabriel-Aundey Risas, and Ben D. Marks (DRC). The authors thank the following Chicago State University students for their assistance with lab work: Korvell Russell, Adaeze Olikagu, Darcae Holmes, Alayjah Harshaw, and Daisy Hernandez. Student training was funded through an IIE Rodman Rockefeller Centennial Fellowship. DJB was supported by the Edward Mallinckrodt, Jr. Foundation, and DJB, GFA, and CJC were supported by NSF DBI 2515340.

## Supplementary Information

Supplementary Data S1: Museum samples screened for coronaviruses and paramyxoviruses.

Fig. S1: Histogram showing coverage of 121 traits across all 194 pteropodid species included in our model.

Fig. S2: Results of hyperparameter tuning, showing the test AUC, sensitivity, and specificity of various models with different combinations of parameters.

Table S1: List of suspected betacoronavirus host species from previously published model (Becker et al. 2022) and number of museum specimens tested for each predicted host species. Table S2: Coverage, definitions, and citations for predictors included in boosted regression analysis, excluding dummy variables for each genus.

Table S3: Hyperparameter values for a generalized boosted regression analysis of pteropodid hosts of paramyxoviruses and resulting trait profiles.

Table S4: Paramyxovirus host species predictions from boosted regression analysis and the number of museum specimens tested for each species.

