## Supplementary Information for "Museum collections and machine learning guide discovery of novel coronaviruses and paramyxoviruses"

**Supplementary Figures and Tables**

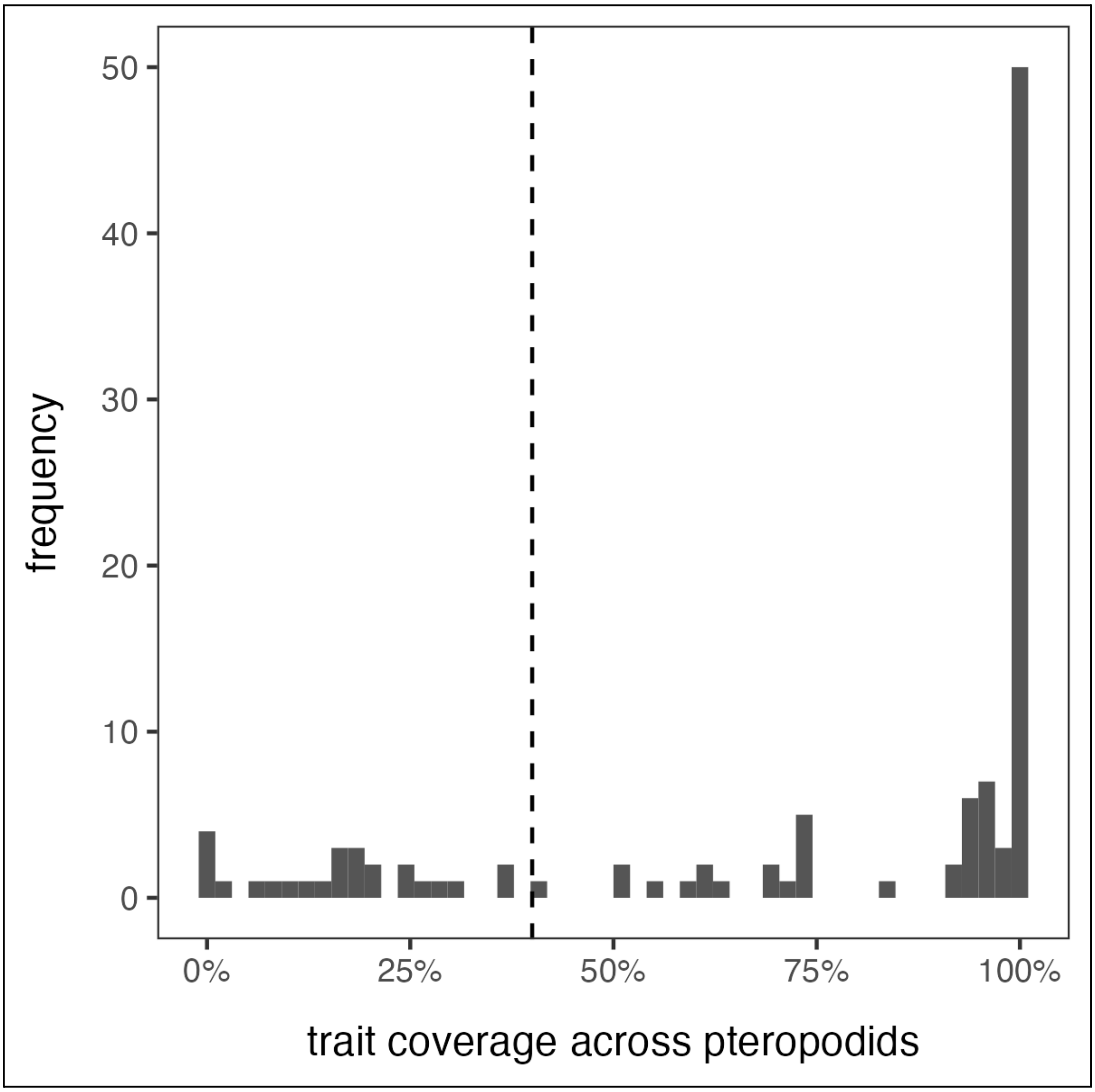

**Fig. S1.** Histogram showing coverage of 121 traits across all 194 pteropodid species included in our model, prior to trimming variables that were completely homogenous across all species and/or missing data for greater than 40% of the species (cutoff indicated with dashed line).

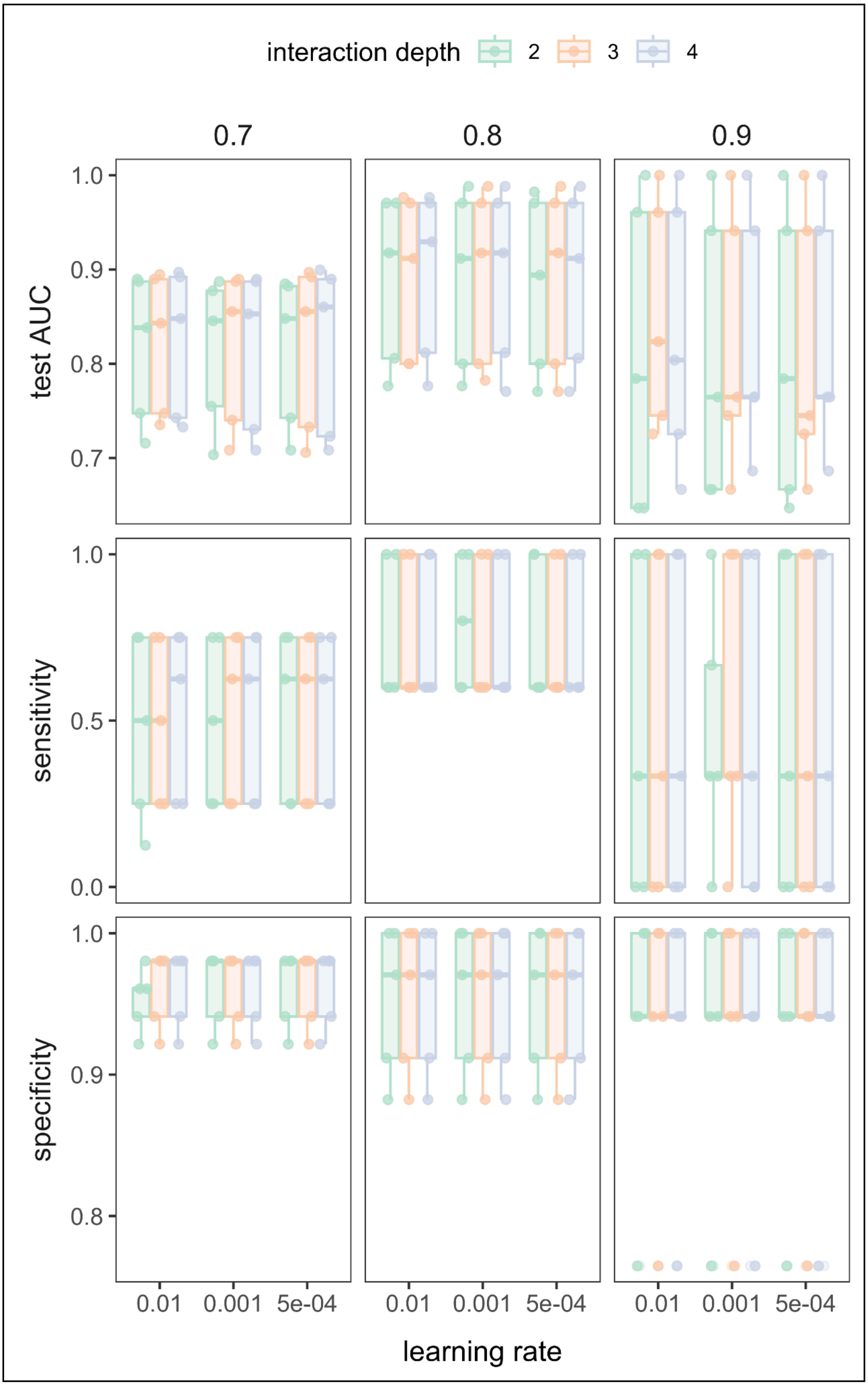

**Fig. S2.** Results of hyperparameter tuning, showing the test AUC, sensitivity, and specificity of various models with different combinations of parameters (interaction depth, learning rate, and percentage of dataset used for training: 70%, 80%, or 90%).

**Table S1.** List of suspected betacoronavirus host species from previously published model (Becker et al. 2022) and number of museum samples tested for each predicted host species.

| Predicted host species | Ensemble.2 | Number of samples tested |
|---|---|---|
| *Haplonycteris fischeri* | 0.19551318 | 92 |
| *Rhinolophus arcuatus* | 0.18658547 | 88 |
| *Chaerephon leucogaster* | 0.16466413 | 64 |
| *Carollia brevicauda* | 0.44939553 | 62 |
| *Glossophaga soricina* | 0.3996464 | 44 |
| *Rhinolophus landeri* | 0.07293978 | 29 |
| *Ptenochirus minor* | 0.27398842 | 27 |
| *Uroderma bilobatum* | 0.4397305 | 27 |
| *Epomophorus wahlbergi* | 0.35274982 | 25 |
| *Alionycteris paucidentata* | 0.48443313 | 24 |
| *Rhinolophus hildebrandtii* | 0.50032908 | 24 |
| *Rhinolophus fumigatus* | 0.04808258 | 22 |
| *Taphozous melanopogon* | 0.21207549 | 22 |
| *Mops sarasinorum* | 0.46699584 | 17 |
| *Harpyionycteris whiteheadi* | 0.19690652 | 16 |
| *Nycticeinops schlieffeni* | 0.05599841 | 15 |
| *Pipistrellus javanicus* | 0.08145241 | 15 |
| *Rhinolophus alcyone* | 0.17034665 | 15 |
| *Rhinolophus inops* | 0.32421467 | 15 |
| *Pteropus hypomelanus* | 0.3293027 | 14 |
| *Sturnira lilium* | 0.35354045 | 14 |
| *Hipposideros obscurus* | 0.44713562 | 13 |
| *Megaerops wetmorei* | 0.14751239 | 11 |
| *Nycteris grandis* | 0.27905654 | 11 |
| *Rousettus lanosus* | 0.23842661 | 11 |
| *Phyllostomus hastatus* | 0.48167335 | 10 |
| *Myotis nigricans* | 0.30855723 | 9 |
| *Rhinolophus virgo* | 0.30411361 | 8 |
| *Casinycteris argynnis* | 0.20356027 | 7 |
| *Myotis riparius* | 0.43457278 | 7 |
| *Molossus rufus* | 0.40932737 | 6 |
| *Philetor brachypterus* | 0.41141958 | 6 |
| *Pteropus pumilus* | 0.22624478 | 6 |
| *Myotis keaysi* | 0.49789229 | 5 |
| *Myotis muricola* | 0.0812209 | 5 |
| *Myotis welwitschii* | 0.26232649 | 5 |
| *Artibeus cinereus* | 0.50662734 | 4 |
| *Epomops dobsonii* | 0.39649016 | 4 |
| *Hypsignathus monstrosus* | 0.3313319 | 4 |
| *Nycteris arge* | 0.36941615 | 4 |
| *Rhinolophus swinnyi* | 0.36653075 | 4 |
| *Scotophilus viridis* | 0.06592144 | 4 |

| | | |
|---|---|---|
| *Epomophorus minor* | 0.26262658 | 3 |
| *Glauconycteris humeralis* | 0.39188126 | 3 |
| *Hipposideros cyclops* | 0.22886078 | 3 |
| *Rhinolophus deckenii* | 0.45344987 | 3 |
| *Rhinolophus simulator* | 0.46615177 | 3 |
| *Scotoecus hindei* | 0.19254345 | 3 |
| *Tadarida brasiliensis* | 0.45297732 | 3 |
| *Hipposideros beatus* | 0.3374091 | 2 |
| *Laephotis wintoni* | 0.31873738 | 2 |
| *Miniopterus tristis* | 0.42449496 | 2 |
| *Myotis bocagii* | 0.07597073 | 2 |
| *Nyctalus leisleri* | 0.0901984 | 2 |
| *Nyctimene rabori* | 0.36679256 | 2 |
| *Pteropus dasymallus* | 0.31397503 | 2 |
| *Rhinolophus maclaudi* | 0.42792758 | 2 |
| *Rhinolophus philippinensis* | 0.221991 | 2 |
| *Eptesicus hottentotus* | 0.31836133 | 1 |
| *Kerivoula argentata* | 0.33100031 | 1 |
| *Kerivoula whiteheadi* | 0.17841583 | 1 |
| *Myonycteris relicta* | 0.29133602 | 1 |
| *Myotis blythii* | 0.22863914 | 1 |
| *Myotis macrotarsus* | 0.23863776 | 1 |
| *Nycteris hispida* | 0.19872946 | 1 |
| *Scotoecus hirundo* | 0.10045996 | 1 |
| *Scotophilus nux* | 0.46209098 | 1 |
| *Acerodon jubatus* | 0.20310657 | 0 |
| *Acerodon leucotis* | 0.37584537 | 0 |
| *Acerodon mackloti* | 0.33338097 | 0 |
| *Aethalops aequalis* | 0.30305896 | 0 |
| *Aethalops alecto* | 0.17954257 | 0 |
| *Arielulus aureocollaris* | 0.30178642 | 0 |
| *Arielulus circumdatus* | 0.13822527 | 0 |
| *Arielulus societatis* | 0.45855507 | 0 |
| *Arielulus torquatus* | 0.37704789 | 0 |
| *Asellia tridens* | 0.1948913 | 0 |
| *Balionycteris maculata* | 0.1394702 | 0 |
| *Barbastella barbastellus* | 0.21950006 | 0 |
| *Barbastella leucomelas* | 0.17430113 | 0 |
| *Chaerephon aloysiisabaudiae* | 0.37783618 | 0 |
| *Chaerephon ansorgei* | 0.34316708 | 0 |
| *Chaerephon bemmeleni* | 0.36763177 | 0 |
| *Chaerephon bivittatus* | 0.41161691 | 0 |
| *Chaerephon chapini* | 0.4828267 | 0 |
| *Chaerephon johorensis* | 0.4992151 | 0 |
| *Chaerephon major* | 0.28523969 | 0 |
| *Chaerephon nigeriae* | 0.29261278 | 0 |
| *Chaerephon russatus* | 0.41693688 | 0 |

| | | |
|---|---|---|
| *Chalinolobus nigrogriseus* | 0.48585933 | 0 |
| *Cheiromeles parvidens* | 0.50246256 | 0 |
| *Cheiromeles torquatus* | 0.3585112 | 0 |
| *Chironax melanocephalus* | 0.14725611 | 0 |
| *Cistugo seabrae* | 0.47719279 | 0 |
| *Coelops frithii* | 0.16898934 | 0 |
| *Coleura afra* | 0.27894663 | 0 |
| *Cynopterus luzoniensis* | 0.27821277 | 0 |
| *Cynopterus minutus* | 0.13465957 | 0 |
| *Cynopterus nusatenggara* | 0.3664413 | 0 |
| *Cynopterus titthaecheilus* | 0.15017815 | 0 |
| *Dobsonia chapmani* | 0.43151864 | 0 |
| *Dobsonia crenulata* | 0.35558202 | 0 |
| *Dobsonia exoleta* | 0.32460793 | 0 |
| *Dobsonia magna* | 0.42776401 | 0 |
| *Dobsonia minor* | 0.35506115 | 0 |
| *Dobsonia peronii* | 0.34488887 | 0 |
| *Dobsonia viridis* | 0.36462844 | 0 |
| *Dyacopterus brooksi* | 0.22527562 | 0 |
| *Emballonura monticola* | 0.38716065 | 0 |
| *Eonycteris major* | 0.20233268 | 0 |
| *Eonycteris robusta* | 0.23668139 | 0 |
| *Epomophorus angolensis* | 0.28119639 | 0 |
| *Epomophorus crypturus* | 0.23529 | 0 |
| *Epomophorus grandis* | 0.32489883 | 0 |
| *Epomophorus minimus* | 0.20939757 | 0 |
| *Eptesicus bobrinskoi* | 0.4429051 | 0 |
| *Eptesicus bottae* | 0.21738853 | 0 |
| *Eptesicus dimissus* | 0.48480563 | 0 |
| *Eptesicus floweri* | 0.3610308 | 0 |
| *Eptesicus fuscus* | 0.35909696 | 0 |
| *Eptesicus gobiensis* | 0.24239754 | 0 |
| *Eptesicus matroka* | 0.42353771 | 0 |
| *Eptesicus nasutus* | 0.25918495 | 0 |
| *Eptesicus pachyotis* | 0.18126307 | 0 |
| *Eptesicus platyops* | 0.18819389 | 0 |
| *Eptesicus tatei* | 0.42726633 | 0 |
| *Eudiscopus denticulus* | 0.28869273 | 0 |
| *Falsistrellus affinis* | 0.15751249 | 0 |
| *Falsistrellus mordax* | 0.3451544 | 0 |
| *Falsistrellus petersi* | 0.22811976 | 0 |
| *Glauconycteris alboguttata* | 0.44895646 | 0 |
| *Glauconycteris beatrix* | 0.25297108 | 0 |
| *Glauconycteris egeria* | 0.39834799 | 0 |
| *Glauconycteris gleni* | 0.40058482 | 0 |
| *Glauconycteris poensis* | 0.14282114 | 0 |
| *Glauconycteris superba* | 0.32929575 | 0 |

| | | |
|---|---|---|
| *Glischropus javanus* | 0.49583535 | 0 |
| *Glischropus tylopus* | 0.09600502 | 0 |
| *Harpiocephalus harpia* | 0.08015329 | 0 |
| *Harpiocephalus mordax* | 0.11270682 | 0 |
| *Harpyionycteris celebensis* | 0.35103623 | 0 |
| *Hesperoptenus blanfordi* | 0.11569942 | 0 |
| *Hesperoptenus tickelli* | 0.13156243 | 0 |
| *Hesperoptenus tomesi* | 0.33421607 | 0 |
| *Hipposideros ater* | 0.11606021 | 0 |
| *Hipposideros bicolor* | 0.29235013 | 0 |
| *Hipposideros cineraceus* | 0.47135237 | 0 |
| *Hipposideros diadema* | 0.43979755 | 0 |
| *Hipposideros dyacorum* | 0.45434796 | 0 |
| *Hipposideros fulvus* | 0.37742472 | 0 |
| *Hipposideros grandis* | 0.31993095 | 0 |
| *Hipposideros halophyllus* | 0.44855957 | 0 |
| *Hipposideros hypophyllus* | 0.34361449 | 0 |
| *Hipposideros jonesi* | 0.35318322 | 0 |
| *Hipposideros lamottei* | 0.50468511 | 0 |
| *Hipposideros lylei* | 0.09678732 | 0 |
| *Hipposideros madurae* | 0.41225175 | 0 |
| *Hipposideros nequam* | 0.43875613 | 0 |
| *Hipposideros orbiculus* | 0.46713618 | 0 |
| *Hipposideros ridleyi* | 0.3743021 | 0 |
| *Hipposideros rotalis* | 0.27512963 | 0 |
| *Hipposideros sorenseni* | 0.46802507 | 0 |
| *Hipposideros turpis* | 0.32076484 | 0 |
| *Hypsugo alaschanicus* | 0.36700985 | 0 |
| *Hypsugo anchietae* | 0.48588085 | 0 |
| *Hypsugo anthonyi* | 0.3793881 | 0 |
| *Hypsugo arabicus* | 0.4088381 | 0 |
| *Hypsugo ariel* | 0.38547157 | 0 |
| *Hypsugo bodenheimeri* | 0.37237169 | 0 |
| *Hypsugo cadornae* | 0.18406542 | 0 |
| *Hypsugo crassulus* | 0.37197654 | 0 |
| *Hypsugo eisentrauti* | 0.36940056 | 0 |
| *Hypsugo imbricatus* | 0.33179516 | 0 |
| *Hypsugo joffrei* | 0.44354994 | 0 |
| *Hypsugo lophurus* | 0.48780737 | 0 |
| *Hypsugo macrotis* | 0.35524199 | 0 |
| *Hypsugo vordermanni* | 0.47993831 | 0 |
| *Kerivoula cuprosa* | 0.38551273 | 0 |
| *Kerivoula flora* | 0.42428268 | 0 |
| *Kerivoula hardwickii* | 0.06925272 | 0 |
| *Kerivoula intermedia* | 0.32099667 | 0 |
| *Kerivoula lanosa* | 0.21151561 | 0 |
| *Kerivoula lenis* | 0.43134915 | 0 |

| | | |
|---|---|---|
| *Kerivoula minuta* | 0.27764606 | 0 |
| *Kerivoula papillosa* | 0.10196111 | 0 |
| *Kerivoula pellucida* | 0.17571296 | 0 |
| *Kerivoula phalaena* | 0.29056098 | 0 |
| *Kerivoula picta* | 0.07933955 | 0 |
| *Kerivoula smithii* | 0.29886523 | 0 |
| *Laephotis botswanae* | 0.41868509 | 0 |
| *Latidens salimalii* | 0.35937488 | 0 |
| *Lavia frons* | 0.32450505 | 0 |
| *Macroglossus sobrinus* | 0.01618042 | 0 |
| *Megaderma lyra* | 0.19086437 | 0 |
| *Megaderma spasma* | 0.17933317 | 0 |
| *Micropteropus intermedius* | 0.2723321 | 0 |
| *Mimetillus moloneyi* | 0.14183201 | 0 |
| *Miniopterus africanus* | 0.30986893 | 0 |
| *Miniopterus medius* | 0.39502997 | 0 |
| *Molossus molossus* | 0.45708699 | 0 |
| *Mops brachypterus* | 0.41086092 | 0 |
| *Mops congicus* | 0.40135724 | 0 |
| *Mops demonstrator* | 0.49701316 | 0 |
| *Mops midas* | 0.30562907 | 0 |
| *Mops mops* | 0.38913182 | 0 |
| *Mops nanulus* | 0.3810738 | 0 |
| *Mops spurrelli* | 0.45163102 | 0 |
| *Mops thersites* | 0.36919736 | 0 |
| *Murina aenea* | 0.33136246 | 0 |
| *Murina aurata* | 0.4931479 | 0 |
| *Murina cyclotis* | 0.08981094 | 0 |
| *Murina florium* | 0.47896721 | 0 |
| *Murina fusca* | 0.2672682 | 0 |
| *Murina grisea* | 0.38417302 | 0 |
| *Murina hilgendorfi* | 0.33365405 | 0 |
| *Murina huttoni* | 0.1809286 | 0 |
| *Murina leucogaster* | 0.16416642 | 0 |
| *Murina puta* | 0.43431169 | 0 |
| *Murina rozendaali* | 0.38853316 | 0 |
| *Murina silvatica* | 0.43170005 | 0 |
| *Murina suilla* | 0.23053498 | 0 |
| *Murina tubinaris* | 0.12781387 | 0 |
| *Myonycteris brachycephala* | 0.39072491 | 0 |
| *Myopterus daubentonii* | 0.35824079 | 0 |
| *Myopterus whitleyi* | 0.45285995 | 0 |
| *Myotis adversus* | 0.16707993 | 0 |
| *Myotis altarium* | 0.06158631 | 0 |
| *Myotis annamiticus* | 0.33878239 | 0 |
| *Myotis annectans* | 0.12667203 | 0 |
| *Myotis ater* | 0.28878809 | 0 |

| | | |
|---|---|---|
| *Myotis bechsteinii* | 0.17290953 | 0 |
| *Myotis bombinus* | 0.3527753 | 0 |
| *Myotis brandtii* | 0.4426578 | 0 |
| *Myotis capaccinii* | 0.44675318 | 0 |
| *Myotis chinensis* | 0.05178499 | 0 |
| *Myotis ciliolabrum* | 0.50761154 | 0 |
| *Myotis csorbai* | 0.31607982 | 0 |
| *Myotis fimbriatus* | 0.23842938 | 0 |
| *Myotis formosus* | 0.08564637 | 0 |
| *Myotis frater* | 0.42104108 | 0 |
| *Myotis gomantongensis* | 0.35693196 | 0 |
| *Myotis goudoti* | 0.41517545 | 0 |
| *Myotis hajastanicus* | 0.3880779 | 0 |
| *Myotis hasseltii* | 0.11764793 | 0 |
| *Myotis hermani* | 0.49414704 | 0 |
| *Myotis ikonnikovi* | 0.32245912 | 0 |
| *Myotis laniger* | 0.12708266 | 0 |
| *Myotis lucifugus* | 0.37404743 | 0 |
| *Myotis moluccarum* | 0.37482842 | 0 |
| *Myotis montivagus* | 0.10402559 | 0 |
| *Myotis morrisi* | 0.32344402 | 0 |
| *Myotis myotis* | 0.22914873 | 0 |
| *Myotis mystacinus* | 0.34938158 | 0 |
| *Myotis nattereri* | 0.23148904 | 0 |
| *Myotis nipalensis* | 0.41734187 | 0 |
| *Myotis occultus* | 0.4904646 | 0 |
| *Myotis oreias* | 0.48816504 | 0 |
| *Myotis oxygnathus* | 0.10203449 | 0 |
| *Myotis pruinosus* | 0.40306317 | 0 |
| *Myotis ridleyi* | 0.25561576 | 0 |
| *Myotis rosseti* | 0.15737882 | 0 |
| *Myotis schaubi* | 0.31267013 | 0 |
| *Myotis scotti* | 0.45204349 | 0 |
| *Myotis septentrionalis* | 0.45742427 | 0 |
| *Myotis sicarius* | 0.26011069 | 0 |
| *Myotis siligorensis* | 0.09293589 | 0 |
| *Myotis tricolor* | 0.23804481 | 0 |
| *Myotis velifer* | 0.4398209 | 0 |
| *Myotis volans* | 0.50954404 | 0 |
| *Neopteryx frosti* | 0.41586109 | 0 |
| *Neoromicia brunneus* | 0.3979622 | 0 |
| *Neoromicia flavescens* | 0.5064773 | 0 |
| *Neoromicia guineensis* | 0.13552339 | 0 |
| *Neoromicia helios* | 0.25934146 | 0 |
| *Neoromicia melckorum* | 0.42725139 | 0 |
| *Neoromicia rendalli* | 0.07092712 | 0 |
| *Neoromicia tenuipinnis* | 0.11127701 | 0 |

| | | |
|---|---|---|
| *Nyctalus aviator* | 0.22292297 | 0 |
| *Nyctalus lasiopterus* | 0.43706984 | 0 |
| *Nyctalus montanus* | 0.25880118 | 0 |
| *Nyctalus plancyi* | 0.4691173 | 0 |
| *Nycteris intermedia* | 0.45029081 | 0 |
| *Nycteris javanica* | 0.48129993 | 0 |
| *Nycteris major* | 0.4200312 | 0 |
| *Nycteris tragata* | 0.29412559 | 0 |
| *Nyctimene aello* | 0.44080709 | 0 |
| *Nyctimene albiventer* | 0.43588329 | 0 |
| *Nyctimene cephalotes* | 0.32271147 | 0 |
| *Nyctimene minutus* | 0.41298438 | 0 |
| *Nyctophilus arnhemensis* | 0.47809963 | 0 |
| *Nyctophilus bifax* | 0.4164495 | 0 |
| *Nyctophilus geoffroyi* | 0.44908793 | 0 |
| *Nyctophilus timoriensis* | 0.49702145 | 0 |
| *Otomops martiensseni* | 0.3907794 | 0 |
| *Otonycteris hemprichii* | 0.2060475 | 0 |
| *Otopteropus cartilagonodus* | 0.29474885 | 0 |
| *Paracoelops megalotis* | 0.36266379 | 0 |
| *Paranyctimene raptor* | 0.46810026 | 0 |
| *Paranyctimene tenax* | 0.46101967 | 0 |
| *Penthetor lucasi* | 0.15109225 | 0 |
| *Phoniscus atrox* | 0.30810259 | 0 |
| *Phoniscus jagorii* | 0.19029576 | 0 |
| *Pipistrellus ceylonicus* | 0.08691568 | 0 |
| *Pipistrellus endoi* | 0.43282039 | 0 |
| *Pipistrellus inexspectatus* | 0.50814699 | 0 |
| *Pipistrellus maderensis* | 0.41060903 | 0 |
| *Pipistrellus nanulus* | 0.2645137 | 0 |
| *Pipistrellus papuanus* | 0.50693949 | 0 |
| *Pipistrellus paterculus* | 0.14873533 | 0 |
| *Pipistrellus rueppellii* | 0.05752272 | 0 |
| *Pipistrellus rusticus* | 0.1263936 | 0 |
| *Pipistrellus stenopterus* | 0.23048792 | 0 |
| *Pipistrellus subflavus* | 0.44844079 | 0 |
| *Pipistrellus westralis* | 0.44663419 | 0 |
| *Plecotus austriacus* | 0.15560912 | 0 |
| *Plecotus taivanus* | 0.49679942 | 0 |
| *Plecotus teneriffae* | 0.47594627 | 0 |
| *Plerotes anchietae* | 0.25538746 | 0 |
| *Pteropus brunneus* | 0.4034434 | 0 |
| *Pteropus caniceps* | 0.41615536 | 0 |
| *Pteropus chrysoproctus* | 0.40948478 | 0 |
| *Pteropus faunulus* | 0.39091346 | 0 |
| *Pteropus griseus* | 0.313766 | 0 |
| *Pteropus intermedius* | 0.13606704 | 0 |

| | | |
|---|---|---|
| *Pteropus leucopterus* | 0.21759303 | 0 |
| *Pteropus lombocensis* | 0.33264041 | 0 |
| *Pteropus loochoensis* | 0.45712179 | 0 |
| *Pteropus macrotis* | 0.41794384 | 0 |
| *Pteropus mariannus* | 0.48368452 | 0 |
| *Pteropus melanopogon* | 0.45778912 | 0 |
| *Pteropus melanotus* | 0.32128746 | 0 |
| *Pteropus niger* | 0.42112798 | 0 |
| *Pteropus ocularis* | 0.42498686 | 0 |
| *Pteropus personatus* | 0.43015853 | 0 |
| *Pteropus pilosus* | 0.42590285 | 0 |
| *Pteropus pohlei* | 0.51037768 | 0 |
| *Pteropus pselaphon* | 0.48788321 | 0 |
| *Pteropus rodricensis* | 0.51108301 | 0 |
| *Pteropus scapulatus* | 0.27901232 | 0 |
| *Pteropus seychellensis* | 0.49591467 | 0 |
| *Pteropus speciosus* | 0.32553876 | 0 |
| *Pteropus subniger* | 0.42803962 | 0 |
| *Pteropus tokudae* | 0.3949762 | 0 |
| *Pteropus vampyrus* | 0.28652479 | 0 |
| *Rhinolophus beddomei* | 0.29008545 | 0 |
| *Rhinolophus bocharicus* | 0.33318634 | 0 |
| *Rhinolophus borneensis* | 0.16164649 | 0 |
| *Rhinolophus canuti* | 0.47419601 | 0 |
| *Rhinolophus celebensis* | 0.30423131 | 0 |
| *Rhinolophus coelophyllus* | 0.38903897 | 0 |
| *Rhinolophus convexus* | 0.26677849 | 0 |
| *Rhinolophus darlingi* | 0.28532343 | 0 |
| *Rhinolophus denti* | 0.25340559 | 0 |
| *Rhinolophus eloquens* | 0.45948827 | 0 |
| *Rhinolophus euryotis* | 0.41534026 | 0 |
| *Rhinolophus formosae* | 0.48019478 | 0 |
| *Rhinolophus guineensis* | 0.30409121 | 0 |
| *Rhinolophus hillorum* | 0.426401 | 0 |
| *Rhinolophus imaizumii* | 0.48990393 | 0 |
| *Rhinolophus keyensis* | 0.47879514 | 0 |
| *Rhinolophus lepidus* | 0.10057796 | 0 |
| *Rhinolophus luctus* | 0.05661354 | 0 |
| *Rhinolophus marshalli* | 0.05246518 | 0 |
| *Rhinolophus mitratus* | 0.2888316 | 0 |
| *Rhinolophus osgoodi* | 0.11367956 | 0 |
| *Rhinolophus paradoxolophus* | 0.10319984 | 0 |
| *Rhinolophus robinsoni* | 0.30403465 | 0 |
| *Rhinolophus rouxii* | 0.42182962 | 0 |
| *Rhinolophus sedulus* | 0.23721316 | 0 |
| *Rhinolophus shortridgei* | 0.24061381 | 0 |
| *Rhinolophus siamensis* | 0.40119697 | 0 |

| | | |
|---|---|---|
| *Rhinolophus silvestris* | 0.50698856 | 0 |
| *Rhinolophus subbadius* | 0.30506757 | 0 |
| *Rhinolophus subrufus* | 0.31082978 | 0 |
| *Rhinolophus trifoliatus* | 0.07822131 | 0 |
| *Rhinolophus yunanensis* | 0.11359972 | 0 |
| *Rhinopoma macinnesi* | 0.38121603 | 0 |
| *Rhinopoma microphyllum* | 0.25965656 | 0 |
| *Rhinopoma muscatellum* | 0.49600279 | 0 |
| *Rousettus bidens* | 0.33884369 | 0 |
| *Rousettus celebensis* | 0.26387458 | 0 |
| *Rousettus linduensis* | 0.43198668 | 0 |
| *Rousettus obliviosus* | 0.43518355 | 0 |
| *Rousettus spinalatus* | 0.20086499 | 0 |
| *Saccolaimus peli* | 0.37818604 | 0 |
| *Saccolaimus saccolaimus* | 0.43486527 | 0 |
| *Scotoecus albigula* | 0.32726402 | 0 |
| *Scotoecus albofuscus* | 0.10266947 | 0 |
| *Scotoecus pallidus* | 0.30738099 | 0 |
| *Scotomanes ornatus* | 0.06766671 | 0 |
| *Scotonycteris ophiodon* | 0.15487988 | 0 |
| *Scotonycteris zenkeri* | 0.11453414 | 0 |
| *Scotophilus celebensis* | 0.43430836 | 0 |
| *Scotophilus collinus* | 0.41026583 | 0 |
| *Scotophilus nigrita* | 0.40822224 | 0 |
| *Scotophilus nucella* | 0.24706568 | 0 |
| *Scotophilus robustus* | 0.43917809 | 0 |
| *Scotorepens greyii* | 0.48455975 | 0 |
| *Scotorepens sanborni* | 0.47227006 | 0 |
| *Scotozous dormeri* | 0.22326266 | 0 |
| *Sphaerias blanfordi* | 0.06328044 | 0 |
| *Styloctenium wallacei* | 0.29955876 | 0 |
| *Tadarida aegyptiaca* | 0.1466381 | 0 |
| *Tadarida insignis* | 0.43650693 | 0 |
| *Tadarida latouchei* | 0.37519804 | 0 |
| *Tadarida ventralis* | 0.47052421 | 0 |
| *Taphozous longimanus* | 0.18485132 | 0 |
| *Taphozous mauritianus* | 0.26101526 | 0 |
| *Taphozous nudiventris* | 0.2909406 | 0 |
| *Thoopterus nigrescens* | 0.30149781 | 0 |
| *Tylonycteris robustula* | 0.43686222 | 0 |
| *Vespadelus caurinus* | 0.50271021 | 0 |

**Table S2.** Coverage, definitions, and citations for predictors included in boosted regression analysis, excluding dummy variables for each genus. Highly skewed predictors were transformed with natural log, as noted below.

| Predictor | Coverage | | Definition | Source |
|---|---|---|---|---|
| | All pteropodids | PMV PCR+ pteropodids | | |
| activity_cycle | 0.96 | 1.00 | Time of day in which the species is most active: Defined as time of the day in which the species carries out most of its activities: 1) nocturnal only; 2) mixed; 3) diurnal only. | Soria et al. 2021 |
| Adult_forearm_mm | 0.98 | 1.00 | Natural log of adult forearm length (in millimeters), defined as the length from the elbow to wrist. | Giannini 2019, supplemented by Soria et al. 2021 |
| Adult_length_mm | 0.92 | 0.96 | Natural log of adult body length (in millimeters), defined as the length from the tip of the nose to the anus or base of the tail. | Giannini 2019, supplemented by Soria et al. 2021 |
| Adult_mass_g | 0.99 | 1.00 | Natural log of adult body mass (in grams). | Giannini 2019, supplemented by Soria et al. 2021 |
| Afrotropical | 0.94 | 1.00 | Whether or not the species is present in the Afrotropical biogeographical region: 1) present; 0) not present. | Soria et al. 2021 |
| altitude_breadth_m | 0.59 | 0.64 | Natural log of the difference between the upper and lower elevation limits at which the species occurs (in meters). | Soria et al. 2021 |
| Australasian | 0.94 | 1.00 | Whether or not the species is present in the Australasian biogeographical region: 1) present; 0) not present. | Soria et al. 2021 |
| cites | 1.00 | 1.00 | Natural log of the number of citations of each bat species in PubMed. | PubMed, extracted on February 22, 2024 with the *easyPubMed* R package |
| det_diet_breadth_n | 0.74 | 0.92 | Number of different dietary categories that each constitute ≥20% of diet (ranging from 1 to 5). | Soria et al. 2021 |
| det_fruit | 0.74 | 0.92 | Percentage (%) of diet composed of fruit or drupes. | Soria et al. 2021 |
| det_nect | 0.74 | 0.92 | Percentage (%) of diet composed of nectar, pollen, plant exudates, or gums. | Soria et al. 2021 |
| det_seed | 0.74 | 0.92 | Percentage (%) of diet composed of seed, maize, nuts, spores, wheat, or grains. | Soria et al. 2021 |
| det_vend | 0.74 | 0.92 | Percentage (%) of diet composed of mammals and birds. | Soria et al. 2021 |

| | | | | |
|---|---|---|---|---|
| dissected_by_mountains | 0.69 | 0.84 | Whether or not the range is dissected by mountains, based on elevation gradients with slopes equal to or higher than 5 degrees: 1) dissected; 0) not dissected. | Soria et al. 2021 |
| dphy_invertebrate | 0.97 | 1.00 | Percentage (%) of the diet composed of invertebrates. | Soria et al. 2021 |
| dphy_plant | 0.97 | 1.00 | Percentage (%) of the diet composed of plants or fungi. | Soria et al. 2021 |
| dphy_vertebrate | 0.97 | 1.00 | Percentage (%) of the diet composed of vertebrates. | Soria et al. 2021 |
| ear_mm | 0.95 | 1.00 | Adult ear length (in millimeters). | Giannini 2019 |
| ed_es | 1.00 | 1.00 | Evolutionary distinctiveness, calculated using a taxonomic backbone (Upham et al. 2019) and the Equal Splits index. | Calculated from Upham et al. 2019 with the *phylogregion* R package |
| ed_fp | 1.00 | 1.00 | Evolutionary distinctiveness, calculated using a taxonomic backbone (Upham et al. 2019) and the Fair Proportion index. | Calculated from Upham et al. 2019 with the *phylogregion* R package |
| habitat_breadth_n | 0.91 | 1.00 | Number of distinct level 1 IUCN habitats suitable for the species (ranging from 1 to 9). | Soria et al. 2021 |
| hibernation_torpor | 0.71 | 0.96 | Whether or not the species experiences hibernation or torpor: 1) yes; 0) no. | Soria et al. 2021, supplemented by Cosentino et al. 2023 |
| hindfoot_mm | 0.84 | 1.00 | Adult hindfoot length (in millimeters). | Giannini 2019 |
| Indomalayan | 0.94 | 1.00 | Whether or not the species is present in the Indomalayan biogeographical region: 1) present; 0) not present. | Soria et al. 2021 |
| island_dwelling | 0.69 | 0.84 | Whether 20% or more of the species breeding range occurs on an island: 1) yes; 0) no. | Soria et al. 2021 |
| litter_size_n | 0.63 | 0.92 | Average number of offspring per litter. | Soria et al. 2021, supplemented by Cosentino et al. 2023 |
| lower_elevation_m | 0.62 | 0.64 | Natural log of lower elevation limit at which the species occurs (in meters). | Soria et al. 2021 |
| max_longevity_d | 0.42 | 0.88 | Maximum reported age at death (in days). | Soria et al. 2021, supplemented by Cosentino et al. 2023 |

| | | | | |
|---|---|---|---|---|
| MeanLat | 0.96 | 1.00 | Mean latitude of the species range (in decimal degrees). | IUCN, extracted on January 25, 2024 |
| MeanLong | 0.96 | 1.00 | Mean longitude of the species range (in decimal degrees). | IUCN, extracted on January 25, 2024 |
| Oceanian | 0.94 | 1.00 | Whether or not the species is present in the Oceanian biogeographical region: 1) present; 0) not present. | Soria et al. 2021 |
| PhyloClust41 | 0.55 | 0.88 | One of five phylogenetic clusters of bat species at 41 million years ago. | Guy et al. 2020 |
| RangeArea | 0.96 | 1.00 | Natural log of species total range area (in square kilometers). | IUCN, extracted on January 25, 2024 |
| Residuals | 0.52 | 0.80 | Residuals from the regression of square root transformed range area on species sympatry. | Guy et al. 2020 |
| StatusM | 1.00 | 1.00 | IUCN status: DD) data deficient; LC) least concern; NT) near threatened; V) vulnerable; E) endangered; CE) critically endangered; EX) extinct. | IUCN, extracted on January 25, 2024 |
| Sympatry | 0.52 | 0.80 | A measure of the number of other species that occur in the same geographic range as the bat species of interest. | Guy et al. 2020 |
| tail_mm | 0.94 | 1.00 | Adult tail length (in millimeters). | Giannini 2019 |
| TrendM | 0.98 | 1.00 | IUCN population trend: S) stable; D) decreasing; I) increasing; U) unknown. | IUCN, extracted on January 25, 2024 |
| upper_elevation_m | 0.62 | 0.76 | Natural log of upper elevation limit at which the species occurs (in meters). | Soria et al. 2021 |
| vcites | 1.00 | 1.00 | Natural log of the number of virus-related citations of each bat species in PubMed (search string including "AND (virus OR viral)"). | PubMed, extracted on February 22, 2024 with the *easyPubMed* R package |

**Table S3:** (A) Hyperparameter values for a generalized boosted regression analysis of pteropodid hosts of paramyxoviruses. (B) Resulting trait profiles from the boosted regression analysis.

| **A** | | |
|---|---|---|
| Error structure | Bernoulli | |
| Train/test fraction | 80/20 | |
| Shrinkage | 0.001 | |
| Interaction depth | 3 | |
| best.iter/n.trees | 2153/3000 | |
| CV folds | 5 | |
| Min obs node | 4 | |
| Train/test AUC | 0.995/0.920 | |
| Mean permuted AUC | 0.626 | |
| Corrected test AUC | 0.794 | |
| **B** | | |
| **Predictor** | **Relative influence** | **Relative standard error** |
| vcites | 40.1016 | 1.50610045 |
| RangeArea | 13.6848 | 0.93455999 |
| cites | 13.512 | 1.15112076 |
| Adult_length_mm | 4.4712 | 0.29246101 |
| ear_mm | 3.4696 | 0.27629422 |
| ed_es | 3.4524 | 0.28313029 |
| altitude_breadth_m | 2.6076 | 0.24544536 |
| MeanLong | 2.3684 | 0.16667601 |
| max_longevity_d | 2.2748 | 0.17066743 |
| litter_size_n | 1.4356 | 0.11226023 |
| PhyloClust41 | 1.308 | 0.13609066 |
| ed_fp | 1.3008 | 0.08810206 |
| MeanLat | 1.2632 | 0.06632677 |
| Sympatry | 1.2232 | 0.09402964 |
| hindfoot_mm | 1.2188 | 0.05502218 |
| upper_elevation_m | 0.9704 | 0.07302621 |
| Residuals | 0.7468 | 0.04671445 |
| TrendM | 0.5296 | 0.02571303 |
| StatusM | 0.5156 | 0.05166262 |
| det_fruit | 0.4816 | 0.05937643 |
| Adult_forearm_mm | 0.4384 | 0.02337862 |
| dissected_by_mountains | 0.4308 | 0.06489201 |
| habitat_breadth_n | 0.3868 | 0.04440991 |
| Adult_mass_g | 0.3208 | 0.0239299 |
| det_nect | 0.2908 | 0.03322108 |
| tail_mm | 0.26 | 0.01845716 |
| island_dwelling | 0.2212 | 0.03599222 |

| | | |
|---|---|---|
| Afrotropical | 0.2168 | 0.03554209 |
| gen_Epomophorus | 0.1952 | 0.02271651 |
| Indomalayan | 0.0896 | 0.01296765 |
| gen_Acerodon | 0.07 | 0.0136504 |
| gen_Myonycteris | 0.0384 | 0.00948121 |
| Australasian | 0.0308 | 0.00475815 |
| det_diet_breadth_n | 0.0256 | 0.00458548 |
| gen_Cynopterus | 0.0252 | 0.00695509 |
| lower_elevation_m | 0.0244 | 0.00529402 |
| gen_Rousettus | 0.0072 | 0.00195959 |
| gen_Pteropus | 0.0012 | 0.00087939 |
| activity_cycle | 0 | 0 |
| det_seed | 0 | 0 |
| det_vend | 0 | 0 |
| dphy_invertebrate | 0 | 0 |
| dphy_plant | 0 | 0 |
| dphy_vertebrate | 0 | 0 |
| gen_Aethalops | 0 | 0 |
| gen_Alionycteris | 0 | 0 |
| gen_Aproteles | 0 | 0 |
| gen_Balionycteris | 0 | 0 |
| gen_Casinycteris | 0 | 0 |
| gen_Chironax | 0 | 0 |
| gen_Desmalopex | 0 | 0 |
| gen_Dobsonia | 0 | 0 |
| gen_Dyacopterus | 0 | 0 |
| gen_Eidolon | 0 | 0 |
| gen_Eonycteris | 0 | 0 |
| gen_Epomops | 0 | 0 |
| gen_Haplonycteris | 0 | 0 |
| gen_Harpyionycteris | 0 | 0 |
| gen_Hypsignathus | 0 | 0 |
| gen_Latidens | 0 | 0 |
| gen_Macroglossus | 0 | 0 |
| gen_Megaerops | 0 | 0 |
| gen_Megaloglossus | 0 | 0 |
| gen_Melonycteris | 0 | 0 |
| gen_Micropteropus | 0 | 0 |
| gen_Mirimiri | 0 | 0 |
| gen_Nanonycteris | 0 | 0 |
| gen_Neopteryx | 0 | 0 |
| gen_Notopteris | 0 | 0 |

| | | |
|---|---|---|
| gen_Nyctimene | 0 | 0 |
| gen_Otopteropus | 0 | 0 |
| gen_Paranyctimene | 0 | 0 |
| gen_Penthetor | 0 | 0 |
| gen_Plerotes | 0 | 0 |
| gen_Ptenochirus | 0 | 0 |
| gen_Pteralopex | 0 | 0 |
| gen_Scotonycteris | 0 | 0 |
| gen_Sphaerias | 0 | 0 |
| gen_Styloctenium | 0 | 0 |
| gen_Syconycteris | 0 | 0 |
| gen_Thoopterus | 0 | 0 |
| hibernation_torpor | 0 | 0 |
| Oceanian | 0 | 0 |

**Table S4:** Paramyxovirus host species predictions from boosted regression analysis and the number of museum specimens tested for each species. Thresholds for positivity were set using a 95% sensitivity threshold with the PresenceAbsence package (Freeman and Moisen 2008). Suspected hosts clear the thresholds for both the PCR+ model (>0.18) and the PCR+ model with mean citation count (>0.10).

| **Species** | **PMV PCR+** | **p(PMV PCR+)** | **p(PMV PCR+), mean citation** | **Host status** | **Number of samples tested** |
|---|---|---|---|---|---|
| *Pteropus_giganteus* | 1 | 0.8677353 | 0.2795034 | true+ | |
| *Pteropus_vampyrus* | 1 | 0.86616027 | 0.2663877 | true+ | |
| *Rousettus_aegyptiacus* | 1 | 0.8226563 | 0.20630486 | true+ | 15 |
| *Eidolon_helvum* | 1 | 0.82246873 | 0.19437492 | true+ | 12 |
| *Epomophorus_gambianus* | 1 | 0.80799957 | 0.27811628 | true+ | |
| *Pteropus_rufus* | 1 | 0.8008099 | 0.19874922 | true+ | |
| *Pteropus_scapulatus* | 1 | 0.79524002 | 0.17908758 | true+ | |
| *Hypsignathus_monstrosus* | 1 | 0.79167322 | 0.19503387 | true+ | 5 |
| *Rousettus_leschenaultii* | 1 | 0.79019942 | 0.15599935 | true+ | |
| *Pteropus_alecto* | 1 | 0.78748233 | 0.1204582 | true+ | |
| *Eonycteris_spelaea* | 1 | 0.77161311 | 0.14195934 | true+ | |
| *Pteropus_lylei* | 1 | 0.75793305 | 0.10294871 | true+ | |
| *Pteropus_hypomelanus* | 1 | 0.74118525 | 0.09773626 | true+ | |
| *Pteropus_poliocephalus* | 1 | 0.74048843 | 0.07311391 | true+ | |
| *Pteropus_conspicillatus* | 1 | 0.73778925 | 0.1376036 | true+ | |
| *Micropteropus_pusillus* | 1 | 0.69648705 | 0.33987798 | true+ | 2 |
| *Eidolon_dupreanum* | 1 | 0.63876002 | 0.10916352 | true+ | |
| *Myonycteris_torquata* | 1 | 0.63836458 | 0.21444989 | true+ | 42 |
| *Rousettus_amplexicaudatus* | 1 | 0.61303884 | 0.19658552 | true+ | |
| *Cynopterus_sphinx* | 0 | 0.57588832 | 0.09049348 | unlikely | |
| *Epomops_franqueti* | 0 | 0.50807607 | 0.15296176 | suspected | 20 |
| *Myonycteris_angolensis* | 1 | 0.39617086 | 0.42307365 | true+ | |
| *Cynopterus_brachyotis* | 0 | 0.39587008 | 0.04340865 | unlikely | 10 |
| *Epomophorus_wahlbergi* | 0 | 0.30589268 | 0.21246707 | suspected | 27 |
| *Megaloglossus_woermanni* | 1 | 0.27376394 | 0.2950229 | true+ | 17 |
| *Rousettus_madagascariensis* | 0 | 0.23032652 | 0.0441434 | unlikely | |
| *Acerodon_celebensis* | 1 | 0.20760845 | 0.20762273 | true+ | |
| *Macroglossus_minimus* | 0 | 0.20368417 | 0.19664011 | suspected | 89 |
| *Thoopterus_nigrescens* | 1 | 0.18542619 | 0.22180873 | true+ | |
| *Pteropus_melanotus* | 1 | 0.1747235 | 0.16995444 | true+ | |
| *Epomophorus_minimus* | 1 | 0.15729065 | 0.19704771 | true+ | |
| *Dobsonia_moluccensis* | 0 | 0.14203787 | 0.1399607 | unlikely | |
| *Pteropus_seychellensis* | 0 | 0.12377793 | 0.12147412 | unlikely | |
| *Scotonycteris_bergmansi* | 0 | 0.11204581 | 0.13561067 | unlikely | 9 |
| *Syconycteris_australis* | 0 | 0.10561252 | 0.12118952 | unlikely | |
| *Acerodon_jubatus* | 0 | 0.10132348 | 0.08837966 | unlikely | |
| *Rousettus_lanosus* | 0 | 0.09776522 | 0.12263413 | unlikely | |
| *Pteropus_tonganus* | 0 | 0.09287367 | 0.09101333 | unlikely | |
| *Thoopterus_suhaniahae* | 0 | 0.08841881 | 0.11088425 | unlikely | |

| | | | | | |
|---|---|---|---|---|---|
| *Micropteropus_intermedius* | 0 | 0.0866139 | 0.11744916 | unlikely | |
| *Epomophorus_labiatus* | 0 | 0.08491185 | 0.08230349 | unlikely | |
| *Epomops_buettikoferi* | 0 | 0.08335962 | 0.07326303 | unlikely | |
| *Epomophorus_crypturus* | 0 | 0.07614268 | 0.10444717 | unlikely | |
| *Nanonycteris_veldkampii* | 0 | 0.07381321 | 0.07382205 | unlikely | |
| *Sphaerias_blanfordi* | 0 | 0.07146814 | 0.08910291 | unlikely | |
| *Megaerops_ecaudatus* | 0 | 0.07010605 | 0.08527862 | unlikely | |
| *Pteropus_macrotis* | 0 | 0.05873294 | 0.10834574 | unlikely | |
| *Megaloglossus_azagnyi* | 0 | 0.05622865 | 0.07259669 | unlikely | |
| *Acerodon_mackloti* | 0 | 0.05047296 | 0.09934184 | unlikely | |
| *Aproteles_bulmerae* | 0 | 0.05037641 | 0.09501413 | unlikely | |
| *Pteropus_dasymallus* | 0 | 0.04899396 | 0.03730692 | unlikely | |
| *Casinycteris_argynnis* | 0 | 0.04731503 | 0.05880664 | unlikely | 14 |
| *Pteropus_aldabrensis* | 0 | 0.04715618 | 0.08065083 | unlikely | |
| *Macroglossus_sobrinus* | 0 | 0.0466393 | 0.05746738 | unlikely | |
| *Epomophorus_angolensis* | 0 | 0.04579557 | 0.06685994 | unlikely | |
| *Nyctimene_albiventer* | 0 | 0.04432437 | 0.06220596 | unlikely | |
| *Pteropus_pselaphon* | 0 | 0.04400306 | 0.07268406 | unlikely | |
| *Pteropus_yapensis* | 0 | 0.04280253 | 0.05924933 | unlikely | |
| *Pteropus_livingstonii* | 0 | 0.0423105 | 0.06534466 | unlikely | |
| *Pteropus_mariannus* | 0 | 0.04212433 | 0.0673405 | unlikely | |
| *Pteropus_admiralitatum* | 0 | 0.04116109 | 0.06378573 | unlikely | |
| *Pteropus_rayneri* | 0 | 0.04089736 | 0.08159853 | unlikely | |
| *Pteropus_pelewensis* | 0 | 0.04014542 | 0.06422082 | unlikely | |
| *Epomophorus_grandis* | 0 | 0.04002114 | 0.05392519 | unlikely | |
| *Balionycteris_maculata* | 0 | 0.03946418 | 0.05096365 | unlikely | |
| *Pteropus_tuberculatus* | 0 | 0.03807295 | 0.05921042 | unlikely | |
| *Chironax_melanocephalus* | 0 | 0.03792403 | 0.05348136 | unlikely | |
| *Ptenochirus_jagori* | 0 | 0.03789498 | 0.03745265 | unlikely | |
| *Pteropus_aruensis* | 0 | 0.03757082 | 0.07054411 | unlikely | |
| *Acerodon_leucotis* | 0 | 0.03735892 | 0.06700967 | unlikely | |
| *Epomops_dobsonii* | 0 | 0.03711808 | 0.06003352 | unlikely | 2 |
| *Megaerops_niphanae* | 0 | 0.03711388 | 0.03645684 | unlikely | |
| *Scotonycteris_occidentalis* | 0 | 0.03703208 | 0.04931299 | unlikely | |
| *Dobsonia_peronii* | 0 | 0.03689188 | 0.05585277 | unlikely | |
| *Pteropus_rennelli* | 0 | 0.03673049 | 0.05864671 | unlikely | |
| *Scotonycteris_zenkeri* | 0 | 0.036338 | 0.04727362 | unlikely | |
| *Dobsonia_chapmani* | 0 | 0.03570949 | 0.05401461 | unlikely | |
| *Cynopterus_titthaecheilus* | 0 | 0.0356518 | 0.03501887 | unlikely | |
| *Syconycteris_hobbit* | 0 | 0.03547701 | 0.05066905 | unlikely | |
| *Pteropus_keyensis* | 0 | 0.03533604 | 0.06829027 | unlikely | |
| *Aethalops_alecto* | 0 | 0.03502712 | 0.0490737 | unlikely | |
| *Myonycteris_brachycephala* | 0 | 0.03456826 | 0.04738893 | unlikely | |
| *Penthetor_lucasi* | 0 | 0.03429916 | 0.03362174 | unlikely | |
| *Pteropus_capistratus* | 0 | 0.03377565 | 0.05339481 | unlikely | |
| *Pteropus_neohibernicus* | 0 | 0.03339167 | 0.05579337 | unlikely | |
| *Myonycteris_relicta* | 0 | 0.03332054 | 0.04523249 | unlikely | |

| | | | | |
|---|---|---|---|---|
| *Pteropus_cognatus* | 0 | 0.03326898 | 0.0541454 | unlikely |
| *Cynopterus_horsfieldii* | 0 | 0.03315304 | 0.04503943 | unlikely |
| *Nyctimene_certans* | 0 | 0.03315192 | 0.04661371 | unlikely |
| *Nyctimene_aello* | 0 | 0.03299114 | 0.04965919 | unlikely |
| *Nyctimene_rabori* | 0 | 0.03265607 | 0.04739927 | unlikely |
| *Haplonycteris_fischeri* | 0 | 0.03251344 | 0.04176015 | unlikely |
| *Casinycteris_campomaanensis* | 0 | 0.03205767 | 0.04271272 | unlikely |
| *Plerotes_anchietae* | 0 | 0.03205497 | 0.04382555 | unlikely |
| *Dobsonia_viridis* | 0 | 0.03204178 | 0.03204476 | unlikely |
| *Pteropus_ocularis* | 0 | 0.03200398 | 0.06100795 | unlikely |
| *Dobsonia_beauforti* | 0 | 0.03195075 | 0.04940375 | unlikely |
| *Dobsonia_anderseni* | 0 | 0.03163043 | 0.05792886 | unlikely |
| *Pteropus_samoensis* | 0 | 0.03155181 | 0.04777148 | unlikely |
| *Nyctimene_cephalotes* | 0 | 0.03127507 | 0.04425707 | unlikely |
| *Pteropus_subniger* | 0 | 0.03122029 | 0.0497408 | unlikely |
| *Paranyctimene_raptor* | 0 | 0.03111927 | 0.04495024 | unlikely |
| *Nyctimene_cyclotis* | 0 | 0.03058304 | 0.0460675 | unlikely |
| *Epomophorus_anselli* | 0 | 0.03047922 | 0.04301174 | unlikely |
| *Pteropus_anetianus* | 0 | 0.03033363 | 0.04633989 | unlikely |
| *Dobsonia_exoleta* | 0 | 0.02966462 | 0.04517683 | unlikely |
| *Pteropus_howensis* | 0 | 0.02952913 | 0.04853427 | unlikely |
| *Pteropus_pohlei* | 0 | 0.02940303 | 0.05376806 | unlikely |
| *Rousettus_obliviosus* | 0 | 0.02929952 | 0.04360902 | unlikely |
| *Paranyctimene_tenax* | 0 | 0.02910986 | 0.04438983 | unlikely |
| *Pteropus_mahaganus* | 0 | 0.02885829 | 0.05010574 | unlikely |
| *Pteropus_argentatus* | 0 | 0.02858421 | 0.03974219 | unlikely |
| *Pteropus_pilosus* | 0 | 0.02845315 | 0.03980397 | unlikely |
| *Pteropus_speciosus* | 0 | 0.02831313 | 0.04791337 | unlikely |
| *Nyctimene_minutus* | 0 | 0.02788212 | 0.03882137 | unlikely |
| *Acerodon_humilis* | 0 | 0.02770944 | 0.04473885 | unlikely |
| *Dyacopterus_spadiceus* | 0 | 0.02758009 | 0.03856491 | unlikely |
| *Pteropus_griseus* | 0 | 0.02724193 | 0.04439648 | unlikely |
| *Nyctimene_robinsoni* | 0 | 0.02712154 | 0.03777338 | unlikely |
| *Pteropus_niger* | 0 | 0.02711879 | 0.04391931 | unlikely |
| *Pteropus_caniceps* | 0 | 0.02701047 | 0.04893927 | unlikely |
| *Aethalops_aequalis* | 0 | 0.02673427 | 0.03760858 | unlikely |
| *Cynopterus_minutus* | 0 | 0.02657577 | 0.03629293 | unlikely |
| *Otopteropus_cartilagonodus* | 0 | 0.02656769 | 0.03694878 | unlikely |
| *Eonycteris_major* | 0 | 0.02652463 | 0.03869222 | unlikely |
| *Dobsonia_minor* | 0 | 0.0263754 | 0.03876372 | unlikely |
| *Dyacopterus_brooksi* | 0 | 0.0261677 | 0.0384821 | unlikely |
| *Nyctimene_vizcaccia* | 0 | 0.02602174 | 0.0382919 | unlikely |
| *Dobsonia_crenulata* | 0 | 0.02583514 | 0.04327377 | unlikely |
| *Pteropus_ornatus* | 0 | 0.0257833 | 0.04080136 | unlikely |
| *Neopteryx_frosti* | 0 | 0.02567699 | 0.04022116 | unlikely |
| *Pteropus_rodricensis* | 0 | 0.02559004 | 0.03768824 | unlikely |
| *Nyctimene_sanctacrucis* | 0 | 0.02540372 | 0.03993774 | unlikely |

| | | | | |
|---|---|---|---|---|
| *Pteropus_voeltzkowi* | 0 | 0.02539054 | 0.04127958 | unlikely |
| *Melonycteris_woodfordi* | 0 | 0.02538647 | 0.03777416 | unlikely |
| *Latidens_salimalii* | 0 | 0.02525987 | 0.03608657 | unlikely |
| *Rousettus_bidens* | 0 | 0.02497085 | 0.0417231 | unlikely |
| *Pteropus_chrysoproctus* | 0 | 0.02494179 | 0.04945538 | unlikely |
| *Harpyionycteris_celebensis* | 0 | 0.02490844 | 0.03733455 | unlikely |
| *Rousettus_celebensis* | 0 | 0.02461468 | 0.0368959 | unlikely |
| *Pteralopex_pulchra* | 0 | 0.02449362 | 0.03839396 | unlikely |
| *Casinycteris_ophiodon* | 0 | 0.02440593 | 0.03288079 | unlikely |
| *Cynopterus_nusatenggara* | 0 | 0.02427816 | 0.03461574 | unlikely |
| *Cynopterus_luzoniensis* | 0 | 0.02408644 | 0.03264202 | unlikely |
| *Nyctimene_masalai* | 0 | 0.02390727 | 0.03424644 | unlikely |
| *Rousettus_spinalatus* | 0 | 0.02377704 | 0.0349796 | unlikely |
| *Pteropus_gilliardorum* | 0 | 0.02377305 | 0.03758767 | unlikely |
| *Pteropus_intermedius* | 0 | 0.02363815 | 0.03938967 | unlikely |
| *Pteropus_brunneus* | 0 | 0.02350739 | 0.03203289 | unlikely |
| *Megaerops_kusnotoi* | 0 | 0.02338001 | 0.03390791 | unlikely |
| *Pteropus_pumilus* | 0 | 0.02330764 | 0.03330922 | unlikely |
| *Alionycteris_paucidentata* | 0 | 0.02321812 | 0.03273408 | unlikely |
| *Pteropus_faunulus* | 0 | 0.02321747 | 0.03673563 | unlikely |
| *Styloctenium_wallacei* | 0 | 0.02314831 | 0.03554505 | unlikely |
| *Desmalopex_leucopterus* | 0 | 0.02308816 | 0.04393734 | unlikely |
| *Syconycteris_carolinae* | 0 | 0.02259137 | 0.03218666 | unlikely |
| *Notopteris_macdonaldi* | 0 | 0.02257646 | 0.03362182 | unlikely |
| *Pteropus_lombocensis* | 0 | 0.02237753 | 0.04174426 | unlikely |
| *Pteralopex_anceps* | 0 | 0.02177641 | 0.03685136 | unlikely |
| *Ptenochirus_minor* | 0 | 0.02175732 | 0.03195422 | unlikely |
| *Nyctimene_keasti* | 0 | 0.02175304 | 0.03203121 | unlikely |
| *Notopteris_neocaledonica* | 0 | 0.02169856 | 0.03167705 | unlikely |
| *Pteropus_loochoensis* | 0 | 0.02151228 | 0.03473628 | unlikely |
| *Dobsonia_emersa* | 0 | 0.02145953 | 0.03812698 | unlikely |
| *Pteropus_melanopogon* | 0 | 0.02138752 | 0.04061368 | unlikely |
| *Pteralopex_atrata* | 0 | 0.02137337 | 0.03673518 | unlikely |
| *Melonycteris_melanops* | 0 | 0.02127268 | 0.03135168 | unlikely |
| *Nyctimene_major* | 0 | 0.02126292 | 0.03229364 | unlikely |
| *Pteropus_ualanus* | 0 | 0.02125265 | 0.03573162 | unlikely |
| *Megaerops_wetmorei* | 0 | 0.02124832 | 0.03034019 | unlikely |
| *Nyctimene_draconilla* | 0 | 0.02119027 | 0.03123666 | unlikely |
| *Rousettus_linduensis* | 0 | 0.02115565 | 0.03150287 | unlikely |
| *Harpyionycteris_whiteheadi* | 0 | 0.02109411 | 0.03223511 | unlikely |
| *Melonycteris_fardoulisi* | 0 | 0.02109333 | 0.03085406 | unlikely |
| *Mirimiri_acrodonta* | 0 | 0.02102923 | 0.03319527 | unlikely |
| *Pteralopex_flanneryi* | 0 | 0.02100216 | 0.03556401 | unlikely |
| *Pteropus_pelagicus* | 0 | 0.02096152 | 0.03169206 | unlikely |
| *Pteropus_vetulus* | 0 | 0.02092462 | 0.03247167 | unlikely |
| *Dobsonia_pannietensis* | 0 | 0.02079504 | 0.03366959 | unlikely |
| *Pteropus_molossinus* | 0 | 0.02065727 | 0.03201679 | unlikely |

| | | | | |
|---|---|---|---|---|
| *Pteropus_tokudae* | 0 | 0.02057625 | 0.03105173 | unlikely |
| *Desmalopex_microleucopterus* | 0 | 0.02050894 | 0.03154096 | unlikely |
| *Pteropus_fundatus* | 0 | 0.02049939 | 0.03178527 | unlikely |
| *Eonycteris_robusta* | 0 | 0.02039133 | 0.03034401 | unlikely |
| *Dobsonia_inermis* | 0 | 0.02014958 | 0.032032 | unlikely |
| *Nyctimene_malaitensis* | 0 | 0.02006223 | 0.02936429 | unlikely |
| *Dyacopterus_rickarti* | 0 | 0.02004063 | 0.02988433 | unlikely |
| *Pteropus_personatus* | 0 | 0.01999243 | 0.03171887 | unlikely |
| *Styloctenium_mindorensis* | 0 | 0.01994434 | 0.03149976 | unlikely |
| *Pteropus_woodfordi* | 0 | 0.01971038 | 0.03074311 | unlikely |
| *Dobsonia_praedatrix* | 0 | 0.0195418 | 0.03116565 | unlikely |
| *Pteropus_nitendiensis* | 0 | 0.01950548 | 0.03083641 | unlikely |
| *Pteropus_temminckii* | 0 | 0.01948811 | 0.03151818 | unlikely |
| *Pteralopex_taki* | 0 | 0.01897644 | 0.03079829 | unlikely |
